# Distinct “driving” versus “modulatory” influences of different visual corticothalamic pathways

**DOI:** 10.1101/2021.03.30.437715

**Authors:** Megan A. Kirchgessner, Alexis D. Franklin, Edward M. Callaway

## Abstract

Higher-order (HO) thalamic nuclei interact extensively with the cerebral cortex and are innervated by excitatory corticothalamic (CT) populations in layers 5 and 6. While these distinct CT projections have long been thought to have different functional influences on the HO thalamus, this has never been directly tested. By optogenetically inactivating different CT populations in the primary visual cortex (V1) of awake mice, we demonstrate that layer 5, but not layer 6, CT projections drive visual responses in the HO visual pulvinar, even while both pathways provide retinotopic, baseline excitation to their thalamic targets. Inactivating the superior colliculus also suppressed visual responses in the pulvinar, demonstrating that cortical layer 5 and subcortical inputs both contribute to HO visual thalamic activity - even at the level of putative single neurons. Altogether, these results indicate a functional division of driver and modulator CT pathways from V1 to the visual thalamus *in vivo*.

## Introduction

The thalamus and its interactions with the cortex are increasingly appreciated as essential for sensory-guided behaviors and complex cognition (Halassa and Sherman, 2019). Yet, the nature of these interactions – how the content and manner of communication through cortico-thalamo-cortical pathways are controlled – has been difficult to decipher. This is especially true as it pertains to “higher-order” (HO) thalamic nuclei, such as the pulvinar in the visual system (also sometimes referred to as “LP” in rodents). The pulvinar has been implicated in synchronizing activity across visual cortical areas to support visual attention (Saalmann et al., 2012; Zhou et al., 2016) and in integrating sensory signals with behavioral context (Blot et al., 2020; Roth et al., 2016). Still, a mechanistic understanding of the complex interactions between the cortex and HO nuclei like the pulvinar has been hindered by incomplete knowledge of the functional impact of their cortical inputs *in vivo*.

While critical questions remain, decades of research into the anatomy and physiology of corticothalamic (CT) circuitry across systems and species have revealed a number of common motifs. For instance, most glutamatergic synapses in the thalamus fall into two major categories, “type 1” and “type 2”, that are characterized by differences in synapse strength, size, number, post-synaptic receptor type, short-term plasticity, and more (Bickford, 2015; Sherman, 2016) (Figure 1A). Their distinctions are noteworthy because these different synapse classes arise from different inputs, and at least in certain thalamic nuclei, they also serve different functions. In the dorsolateral geniculate nucleus (dLGN) - the first-order (FO) thalamic nucleus in the visual system – retinal ganglion cells make type 2 synapses and are characterized as “drivers” because they “drive” visual activity in the dLGN (Cleland et al., 1971; Sherman and Guillery, 1996, 1998, 2002; Usrey et al., 1999). In contrast, layer 6 corticothalamic neurons (L6CTs) in the primary visual cortex (V1) make type 1 synapses and are classified as “modulators” because they are not responsible for the dLGN’s visual responses but instead exert more subtle, “modulatory” influences (Briggs and Usrey, 2008; Sherman and Guillery, 1996, 1998, 2002). These classes are not specific to the dLGN; FO nuclei in other sensory systems also receive peripheral “driving” and cortical L6CT “modulatory” inputs (Rouiller and Welker, 2000; Sherman, 2016; Sherman and Guillery, 1998, 2002).

**Figure 1.**
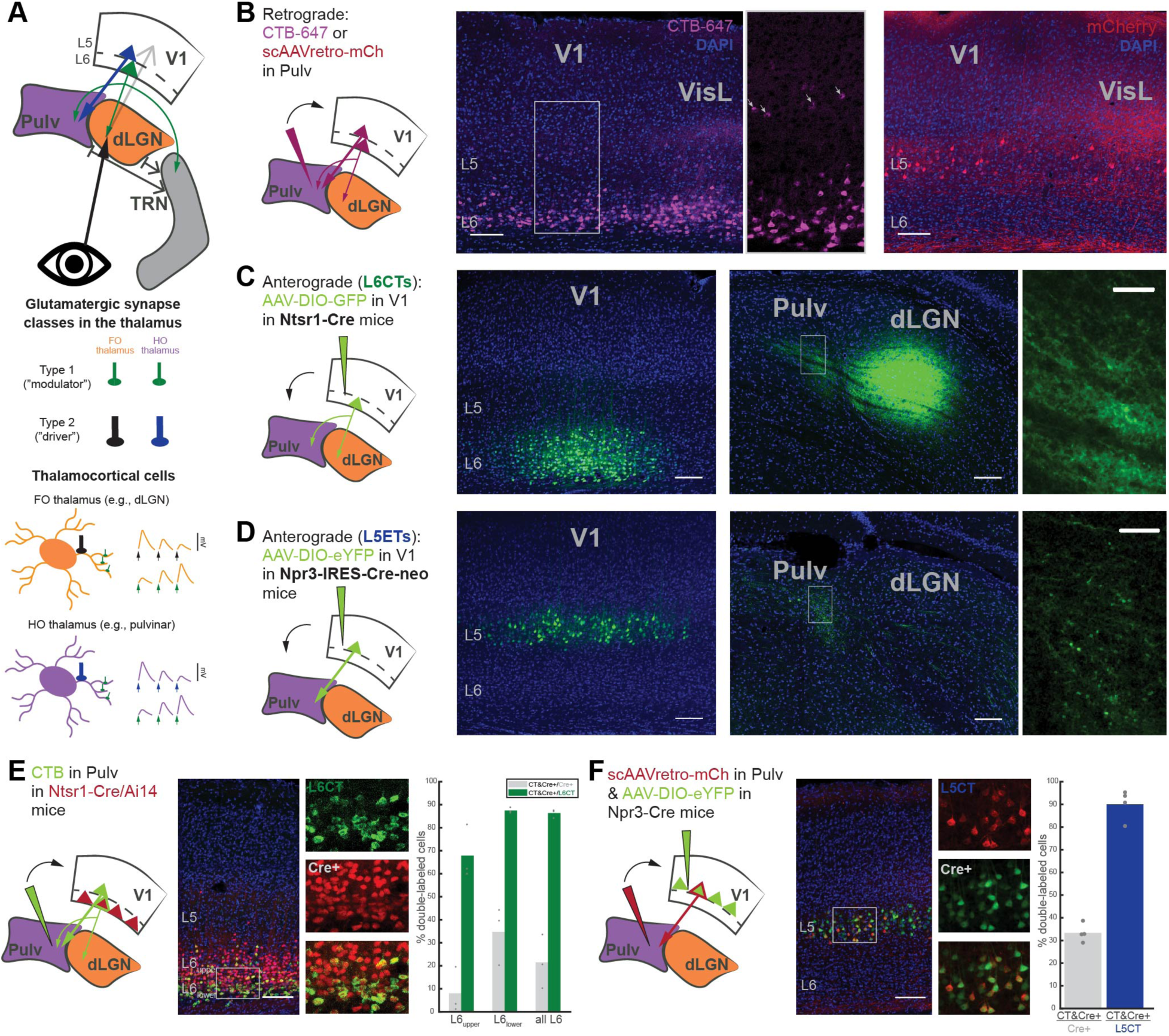
The mouse pulvinar receives V1 input from two distinct corticothalamic (CT) populations that are selectively targeted by different Cre mouse lines. (A) Schematic of the visual CT circuit and different glutamatergic synapse classes. Bottom: cartoons of thalamocortical neurons in first-order (FO; e.g., dLGN) and higher-order (HO; e.g., pulvinar) thalamic nuclei and their distinct inputs, which exhibit different anatomical and physiological properties (e.g., size, number and location of boutons, left; and short-term plasticity dynamics, right). (B) Retrograde tracing from the pulvinar (left) with CTB-647 (middle) or a self-complimenting (sc)AAVretro-mCherry (right), which exhibit different propensities for labeling L6CTs versus L5CTs, respectively. (C) AAV injection of Cre-dependent GFP into V1 of Ntsr1-Cre transgenic mice for L6CT labeling. As expected from the known circuitry (see a), L6CT axons are visible in the dLGN and pulvinar with small, diffuse terminals (right; higher magnification view of boxed region). (D) AAV injection of Cre-dependent eYFP into V1 of Npr3-IRES-Cre-neo mice for labeling L5 extratelencephalic cells (L5ETs). L5 axons are only present in the pulvinar (not dLGN) and exhibit larger and sparser terminals compared to those from L6 (right). (E) Retrograde labeling (using CTB) from the pulvinar in Ntsr1-Cre/Ai14 mice. Individual and composite channel images are from the boxed region. Right: Quantification of double-labeled cells out of all tdTomato+ cells (Cre+) and out of all CTB+ (L6CT) cells in L6_upper_, L6_lower_, and all of L6. n=3 mice. (F) Retrograde labeling (using scAAVretro-mCherry) from the pulvinar in Npr3-Cre mice with AAV injection of Cre-dependent eYFP into V1. Right: quantification of double-labeled cells out of all eYFP+ cells (Cre+) and out of all mCherry+ cells (L5CT; n=4 mice). All scale bars are 100µm for full images and 25µm for higher-magnification images (C, D). Bar graphs depict means across animals, and points are individual animals.

Meanwhile, the functional contributions of different inputs to HO nuclei like the pulvinar is much less clear. HO nuclei (like FO) are also innervated by L6CTs with type 1, “modulator” synapses, but uniquely receive an additional cortical input from layer 5 neurons (L5CTs) with type 2, “driver”-like characteristics (Bourassa and Deschênes, 1995; Li et al., 2003b, 2003a; Mathers, 1972; Reichova and Sherman, 2004; Rockland, 1996) (Figure 1A). L5CTs are thus frequently referred to as “drivers” and L6CTs as “modulators” of HO nuclei (Sherman, 2016; Sherman and Guillery, 2002), but whether these parallel CT projection pathways are functionally distinct has never been directly tested (Bickford, 2015). Experiments demonstrating V1 driving influences on the pulvinar, as well as of the cortex on other HO thalamic nuclei, have relied on inferences following complete (chronic or transient) cortical inactivation (Beltramo and Scanziani, 2019; Bender, 1983; Bennett et al., 2019; Casanova et al., 1997; Diamond et al., 1992; Mease et al., 2016). However, these non-specific approaches would be expected to affect both L5 and L6 CT pathways, as well as activity in other cortical areas and subcortical structures. Thus, the functional consequences of type 1 (L6CT) versus type 2 (L5CT) inputs to the HO thalamus are not known and might differ considerably from the FO thalamus. For instance, type 1 compared to type 2 boutons are significantly more prevalent in HO than in FO thalamic nuclei (Horn and Sherman, 2007; Horn et al., 2000; Wang et al., 2002), suggesting that L6CTs could play a more prominent role in shaping visual responses in the pulvinar than in the dLGN. Therefore, the longstanding question of whether L5 versus L6 CT projections from V1 to the pulvinar are dissociable “driving” versus “modulatory” pathways has yet to be answered.

Another distinguishing feature of HO nuclei is their diversity of excitatory input sources. The rodent pulvinar, for instance, receives additional L5 and L6 CT projections from extrastriate cortical areas (Blot et al., 2020; Roth et al., 2016; Scholl et al., 2020; Souza et al., 2020), as well as subcortical excitation from the superior colliculus (SC) (Zhou et al., 2017). Notably, these SC projections display a mix of “type 1” and “type 2” characteristics (Bickford, 2015) and have even been shown to drive activity in the caudomedial subdivision of the rodent pulvinar (cmPulv) (Beltramo and Scanziani, 2019; Bennett et al., 2019). In the lateral subdivision that receives both cortical and SC input (lPulv) (Zhou et al., 2017), it is unclear whether SC projections are still drivers or how (and if) they might interact with cortical afferents to shape visual responses. These diverse cortical and subcortical projections could suggest a role for the pulvinar as an “information hub” for intersecting pathways (feedforward and feedback, visual and more), putatively involved in multisensory and sensorimotor integration and conveying diverse signals to the cortex (Blot et al., 2020; Froesel et al., 2021; Grieve et al., 2000; Jaramillo et al., 2019; Roth et al., 2016). Yet how these different inputs – L5CT versus L6CT, cortical versus subcortical – interact in the pulvinar *in vivo* is not presently known.

Here, we utilize a variety of viral and transgenic approaches to selectively inactivate L5 versus L6 CT projections from V1 to the pulvinar and dLGN in awake mice and assess their contributions to thalamic activity and visual responses. We find that inactivating either L5CTs or L6CTs in V1 can reduce spontaneous activity in retinotopically aligned thalamic neurons, yet only L5CT inactivation profoundly suppresses visual activity and receptive field properties in the pulvinar. Dual wavelength optogenetics experiments confirmed that individual pulvinar neurons’ visual activity is suppressed by inactivating L5, but not L6, CT projections from V1. Thus, L6CTs in V1 are not required for visual responses in the thalamus and are therefore modulatory, whereas L5CTs provide visual drive to some, though not all, pulvinar neurons. We also investigated the effects of SC inactivation combined with V1 L5CT inactivation and found a variety of neurons whose visual responses are driven by V1 L5CTs, by the SC, or even by both, as well as neurons with undetermined driving sources. Altogether, our findings confirm a longstanding but untested hypothesis regarding the dissociable functions of distinct CT projection populations, and also highlight the role of the rodent HO visual thalamus in integrating inputs from both cortical and subcortical sources as opposed to a strictly feed-forward trans-thalamic pathway.

## Results

### Selective optogenetic inactivation of L6 vs. L5 corticothalamic pathways using different Cre transgenic mouse lines

The mouse pulvinar receives cortical input from two excitatory cell types in V1: L6CTs and L5CTs. Consistent with prior studies (Blot et al., 2020; Roth et al., 2016; Scholl et al., 2020; Souza et al., 2020), we confirmed that injections of a cholera toxin subunit B (CTB) retrograde tracer into the pulvinar labeled CT neurons in V1 L6 – primarily in the lower half of L6 - and in L5, as well as in L6 and L5 of surrounding extrastriate cortical areas (Figure 1B). We also observed more prominent L5 labeling when using a self-complimenting retrograde AAV (scAAVretro) injected into the pulvinar to retrogradely express a fluorophore without infecting L6CTs (which is characteristic of AAVretro (Tervo et al., 2016); Figure 1B).

To target each of these populations in V1 for specific optogenetic manipulations, we utilized two Cre driver lines. For L6CTs, we used the Ntsr1-Cre GN220 BAC transgenic mouse line (Gong et al., 2007), which labels dLGN-projecting L6CTs with near-perfect efficiency and specificity (Bortone et al., 2014; Olsen et al., 2012) and which we previously used to optogenetically stimulate L6CT projections to the dLGN and pulvinar (Kirchgessner et al., 2020). As expected, following injection of Cre-dependent AAV in V1, type 1 axons were observed in both the pulvinar and dLGN (Figure 1C). However, not all V1 L6CTs also project to the pulvinar. L6CTs in V1 that were retrogradely labeled from the pulvinar were largely restricted to lower L6 (Blot et al., 2020; Roth et al., 2016; Souza et al., 2020) (Figures 1B and 1E) and were still a relative minority of all L6CT cells (34.71% in lower L6, 21.5% in all of L6), although these proportions are likely underestimates as a consequence of restricting our injections to avoid the dLGN. Nevertheless, nearly all (87.49% in lower L6, 86.31% in all of L6) pulvinar-projecting L6CTs were labeled (Figure 1E), thus validating the use of the Ntsr1-Cre line for reliably targeting the L6CT projection pathway for optogenetic manipulation.

To inactivate the L5 CT pathway, we characterized and used a new knock-in transgenic mouse line, Npr3-IRES-Cre-neo (Daigle et al., 2018), that showed considerable efficiency and specificity for L5 extratelencephalic neurons (L5ETs; Figures 1D and 1F). In addition to their projections to HO thalamic nuclei, L5ETs can also project to other subcortical structures, such as the SC, and not all L5ETs have collaterals to the thalamus (i.e., a subset of L5ETs are L5CTs; Figures S1A and S1F-H). This line is more specific than the popular Rbp4-Cre line, which labels not only L5ETs, but also L5 cortico-cortical (CC) neurons (Harris et al., 2019; Tasic et al., 2018) (Figures S1A and S1B) that are involved in direct inter- and intra-cortical signaling and thus could confound interpretations about the L5-specific CT pathway from V1. Anterograde tracing from V1 in Npr3-Cre mice yielded axons specifically in the pulvinar, but not the dLGN, with relatively large and sparse terminals – as expected from the known anatomy of L5 CT projections (Bourassa and Deschênes, 1995) (Figure 1D). Indeed, we found that virtually all (90.08%) L5 neurons retrogradely-labeled from the pulvinar (L5CTs) co-expressed a Cre-dependent fluorophore following AAV injection (Figure 1F) and were completely non-overlapping with retrogradely labeled CC neurons (Figures S1C-E). A third (33.33%) of these Cre+ neurons projected to the pulvinar (i.e., ∼1/3 of L5ETs are L5CTs), which is expected given the heterogeneity of L5ETs’ subcortical projection targets (Figure S1). Therefore, the Npr3-Cre line allows for privileged access to L5ET neurons, including the putative “driving” CT pathway from V1 to the pulvinar.

Having verified the specificity of these mouse lines, we preceded to inject Ntsr1-Cre and Npr3-Cre mice with an AAV into V1 to express halorhodopsin in L6CTs or L5ETs, respectively (Figures 2A-C). We then used high-density silicon microprobes (Du et al., 2011) to record extracellular single-unit activity from V1 in awake, headfixed mice viewing full-field square-wave drifting gratings to verify the efficacy and specificity of optogenetic inactivation. Current source density analysis was used to determine the laminar location of each recorded unit in an unbiased manner (Figure 2D). Strikingly, red light used to stimulate halorhodopsin in each of these mouse lines resulted in potent and layer-specific inactivation (Figures 2E and 2F). Because each of these lines only labels subpopulations of excitatory neurons within a layer (L6CT or L5ET), not all regular-spiking (i.e., excitatory) units were silenced, as expected. At least 47/223 (21.08%) and 64/108 (59.26%) regular-spiking units in layers 6 and 5, respectively, putatively expressed halorhodopsin (>50% reduction in visually evoked firing rates, Figures 2G and 2H; example “inactivated” units in Figures 2I and 2J). While those in L6 were fewer than would have been expected given the known proportions of excitatory cell types in L6 (Olsen et al., 2012), L6CTs are likely undersampled in our recordings as they are relatively small pyramidal cells with exceptionally sparse and tuned activity (Vélez-Fort et al., 2014), compared to L5ETs which are large cells with high (∼10Hz) spontaneous firing rates and less tuning selectivity (Williamson and Polley, 2019) (Figure S2C). The fact that we observed increased activity across other layers upon L6CT inactivation, which has been previously reported in mouse V1 (Olsen et al., 2012), provides additional evidence that our suppression was effective (Figure S2A). Altogether, we have identified Cre-driver mouse lines, including a L5ET-specific line, which allow for layer- and cell type-specific optogenetic inactivation of distinct CT projection populations.

**Figure 2.**
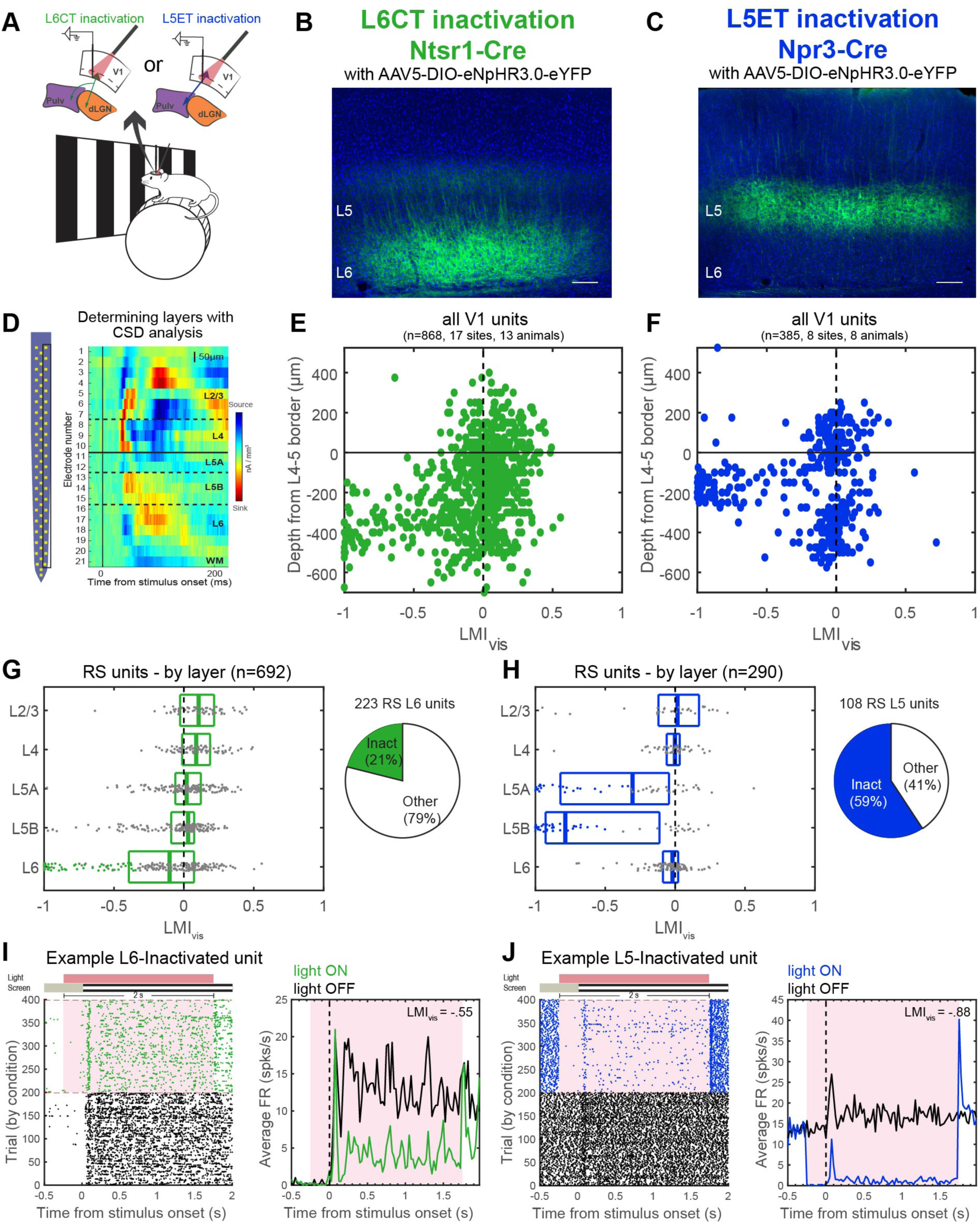
Selective inactivation of L6CTs or L5ETs in the primary visual cortex (V1) of awake mice. (A) Experiment schematic. Mice were awake and headfixed on a wheel and viewing full-field drifting gratings during laminar V1 recordings. (B-C) An AAV encoding Cre-dependent halorhodopsin (eNpHR3.0-eYFP) was injected into V1 in Ntsr1-Cre (L6CTs, B) or Npr3-Cre (L5ETs, C) mice. (D) Left: schematic of a 64-channel silicon probe typically used for V1 recordings. Right: current source density plot for the right column of channels in an example recording. Layers corresponding to each channel’s location were defined by an initial current sink in L4 and a delayed sink in L5B. (E-F) Light modulation index ((FR_lightON_-FR_lightOFF_)/(FR_lightON_+FR_lightOFF_)) from visual trials (LMI_vis_) for all units by depth relative to the L4-5 border if L6CT (E) and L5ET (F) inactivation experiments. (G) Left: LMI_vis_ by layer for regular-spiking units (RS) in L6CT inactivation experiments. Boxplots display the median and quartiles, and putative inactivated units (L6 RS and LMI_vis_ <-.33) are colored dots. Right: proportion of all L6 RS units that were putatively directly inactivated. (H) Same as (G) but for L5ET inactivation experiments. (I) An example inactivated L6 unit. Left: raster plot of all trials organized by condition (green=light ON trials for L6CT inactivation). Right: peristimulus time histogram (PSTH) of average firing rates across visual light OFF (black) or light ON (green) trials. Red shading corresponds to the period of light stimulation. (J) Same as (I) but for an example inactivated L5 unit in L5ET inactivation experiments.

### L6 vs. L5 CT pathways differ in their effects on visually evoked activity in the visual thalamus

We next turned to the thalamus to determine how inactivating L6 versus L5 CT projections from V1 affected activity and visual processing in the lateral subdivision of the pulvinar (lPulv), which receives direct V1 input, in awake mice (Figure 3). Because V1 L6CT feedback to dLGN constitutes a known “modulatory” pathway (Sherman and Guillery, 1998), we also probed the effects of L6CT inactivation on activity in the dLGN. There, L6CT inactivation had disparate effects on spontaneous versus visually evoked activity. In line with another recent study in awake mice (Born et al., 2020), we observed considerable suppression of spontaneous activity in a number of units (Figure 3C) and across the population as a whole (Figures 3D and S3A). However, the presence of a visual stimulus greatly reduced – and in most cases eliminated – any suppression of dLGN activity induced by L6CT inactivation (Figures 3C and 3D), and even subtly increased the temporally modulated (“F1”) response (Figure S3A). Suppressed spontaneous and enhanced F1 responses were also observed in an additional set of experiments in which V1 was inactivated non-specifically (Figure S3D). Because of the distinct effects on spontaneous versus visually evoked activity, “suppressed” cells here and throughout this study (unless otherwise indicated) are those whose spontaneous activity was significantly reduced by the optogenetic manipulation (see Methods). In lPulv, L6CT inactivation did not suppress visually evoked activity, similar to the dLGN. There were also fewer instances of reduced spontaneous activity in lPulv than in the dLGN (although some units exhibited transient suppression in the first ∼200ms of light onset; e.g., Figure 3E), Therefore, L6CT innervation is not required for visual responses in the dLGN or pulvinar, consistent with a fundamentally modulatory role for these projections in both FO and HO nuclei. They can, however, contribute to baseline activity, particularly in the dLGN.

**Figure 3.**
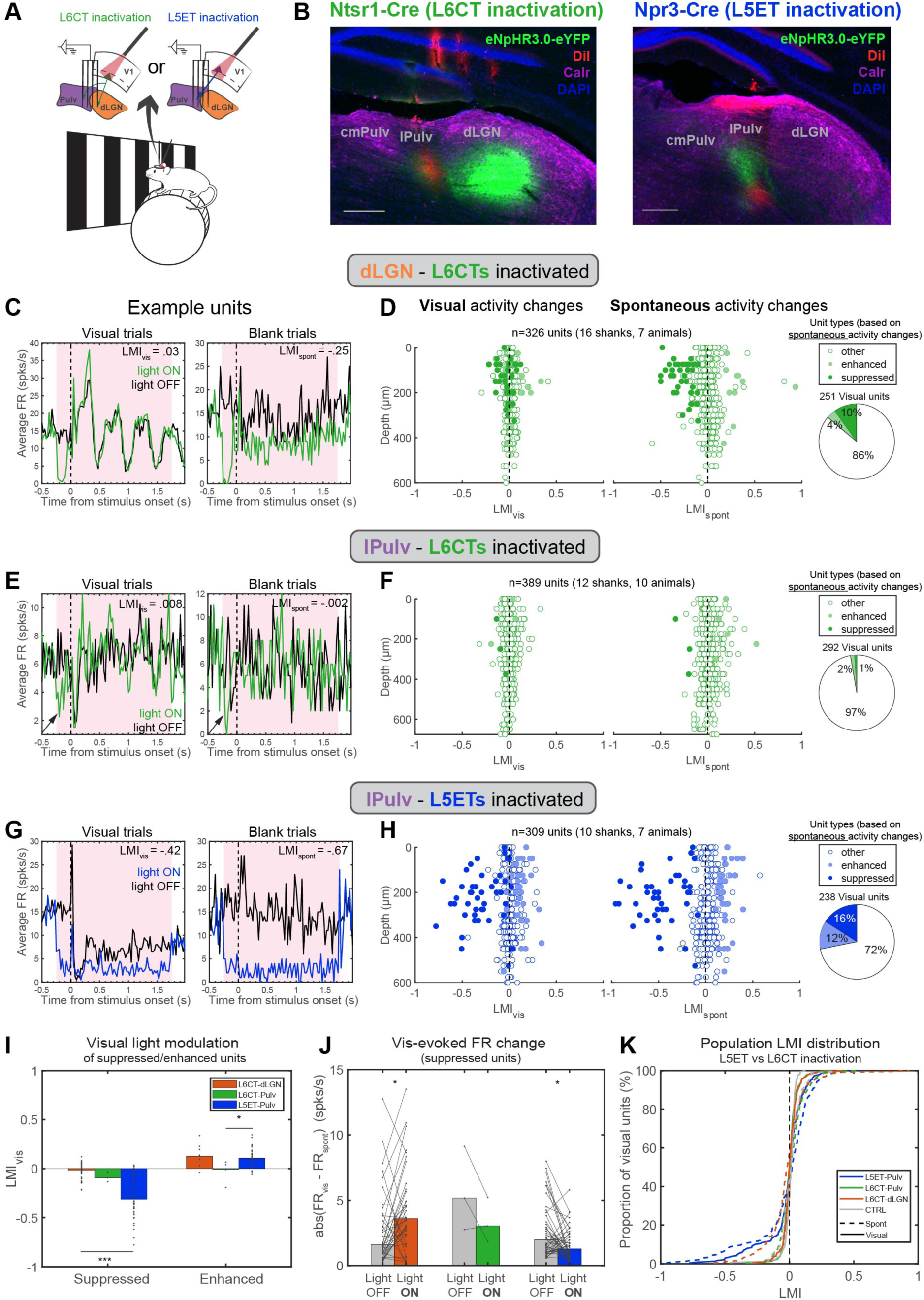
L6CT vs. L5ET inactivation has distinct effects on spontaneous and visually evoked activity in the dLGN and lateral pulvinar (lPulv). (A) Experiment schematic - thalamus recordings in the same animals as in Figure 2. (B) Histological verification of lPulv recording locations (probe shanks coated in DiI, red) in L6CT (left) and L5ET (right) inactivation experiments. Immunohistochemical staining for calretinin (purple) delineates borders between cmPulv (Calr+ cell bodies), lPulv (Calr-), and dLGN (Calr+ axons). Scale bars = 200µm. (C) An example dLGN unit whose spontaneous (right), but not visually evoked (left), activity is suppressed by L6CT inactivation. Plots are PSTHs of average firing rates (FRs, spks/s) across all visual (left) and blank (right) trials under different L6CT inactivation conditions. (D) LMIs of visually evoked (left) and spontaneous (right) activity from all recorded units in dLGN, by depth. Units are colored according to whether their spontaneous activity was significantly suppressed or enhanced (or unaffected) by L6CT inactivation. Far-right: proportion of visually responsive units suppressed, enhanced, or non-modulated (“other”). (E) An example lPulv unit unaffected by L6CT inactivation except at the onset of inactivation in both visual and blank trials (indicated by arrows). (F) Same as (D) but for lPulv units. (G) An example lPulv unit whose visually evoked and spontaneous activity is suppressed by L5ET inactivation. (H) Same as D) but for lPulv units in L5ET inactivation experiments. (I) Median visual LMIs for suppressed and enhanced dLGN and lPulv units under L6CT vs. L5ET inactivation conditions. p<.0001 for L5ET-Pulv vs. L6CT-dLGN, suppressed units, and p=.044 for L5ET-Pulv vs L6CT-Pulv, enhanced units; all other comparisons n.s. (Kruskal-Wallis non-parametric test with the Dunn–Šidák post-hoc test for multiple comparisons). (J) Average visually evoked FR changes (abs(FR_vis_-FR_spont_)) in Light-OFF vs. Light-ON trials for dLGN units in L6CT inactivation experiments (p=0.016), lPulv units in L6CT inactivation experiments (p=0.5), and lPulv units in L5ET inactivation experiments (p=0.012), Wilcoxon signed-rank tests. (K) Distribution of visual and spontaneous LMIs for visually responsive dLGN units in L6CT inactivation experiments (orange), lPulv units in L6CT (green) and L5ET (blue) inactivation experiments, and combined lPulv and dLGN units in control experiments (grey).

V1 L5ET inactivation, on the other hand, had substantial effects not only on baseline activity, but also on visual responses in lPulv neurons (e.g., Figure 3G). Units whose spontaneous activity was significantly suppressed by L5ET inactivation also exhibited considerably reduced activity in the presence of the drifting grating stimulus (Figure 3H). Consequently, the magnitude of optogenetic modulation of visual responses (visual light modulation index, LMI_vis_) was greater in L5ET-suppressed lPulv neurons compared to L6CT-suppressed neurons in either the dLGN or the few in lPulv (Figure 3I). Considered another way, L5ET inactivation reduced the effect of the visual stimulus on firing rates (magnitude of difference between visual and spontaneous firing rates, Figure 3J) in suppressed lPulv units, suggesting that some degree of visual information is conveyed to these pulvinar neurons through the L5 CT pathway. In contrast, visually induced activity changes somewhat increased in L6CT-suppressed dLGN units, as a consequence of their reduced baseline but unchanged visually evoked firing rates (Figure 3J). These differences cannot be attributed to differences in firing mode induced by L6CT versus L5ET inactivation because we observed increased bursting activity in both cases (Figures S3A-C). They are also not exclusive to lPulv, as we also observed suppressed visual activity during L5ET but not L6CT inactivation in the rostromedial pulvinar (rmPulv), which also receives direct V1 input (Figures S3E-H); we therefore combine data from lPulv and rmPulv (collectively referred to as “pulvinar”) for all further analyses unless indicated otherwise. Altogether, we have observed a striking dissociation between the effects of L6CT versus L5ET inactivation on visual activity in the thalamus. This distinction is consistent with the predicted “driving” function of the V1 L5 CT pathway, in contrast to the “modulatory” L6 CT pathway (Sherman and Guillery, 1998).

Nevertheless, it is notable that even the “driving” effects of L5ET inactivation were observed in a minority of visually responsive pulvinar units (Figures 3H and 3K). This contrasts with some expectations from proposed models of L5 “drivers” mediating a trans-thalamic feedforward pathway from primary to higher-order cortical areas (Kato, 1990; Rouiller and Welker, 2000; Sherman and Guillery, 1996, 2002; Theyel et al., 2010). We therefore sought to gain further insight into the functional organization of these distinct CT “driving” and “modulatory” pathways.

### Retinotopic organization of both L6 and L5 excitatory CT projections to the dLGN and pulvinar

The pulvinar is topographically organized – both in its representations of visual space (i.e., retinotopy; Allen et al., 2016; Baldwin et al., 2017; Bennett et al., 2019; Roth et al., 2016) and its connections with visual cortical areas (Juavinett et al., 2020; Tohmi et al., 2014). V1 projections to the pulvinar (Bennett et al., 2019), as well as pulvinar projections back to V1 (Roth et al., 2016), are also coarsely retinotopic. Even though we limited our pulvinar analyses to recordings in which the recording shank passed through CT terminals from V1, nearby pulvinar neurons can have RFs separated by more than 20° (Bennett et al., 2019; Chalupa et al., 1983). Thus, pulvinar neurons that were not modulated by L5ET suppression might simply have not been retinotopically aligned to the cortical inactivation area, causing us to underestimate the “driving” influence of V1 L5 CT projections on the pulvinar. Additionally, given recent findings of spatially organized effects of L6CT activation (Born et al., 2020), we wondered whether retinotopy might also explain some of the effects of L6CT inactivation on spontaneous activity that we observed in the dLGN, and more modestly in the pulvinar.

To identify receptive field (RF) locations of V1 and thalamic neurons, we utilized a sparse noise stimulus protocol (Methods). This allowed us to identify both “on” and “off” subfields (in response to luminance increases and decreases) and to coordinate L6CT or L5ET inactivation with the visual stimulation (Figure 4A). An example L6CT inactivation experiment, in which V1, dLGN and lPulv were all recorded from simultaneously, is depicted in Figure 4B; and a L5ET inactivation experiment with consecutive V1 and lPulv recordings is shown in Figure 4C. As expected, V1 neurons recorded across layers were all retinotopically aligned, whereas thalamus penetrations yielded units whose RFs moved along the elevation and/or azimuth axes according to recording depth. For units whose RFs could be confidently identified (see Methods for criteria), their RF information was then related to the effects of optogenetic inactivation on their responses to drifting grating stimuli (from Figure 3).

**Figure 4.**
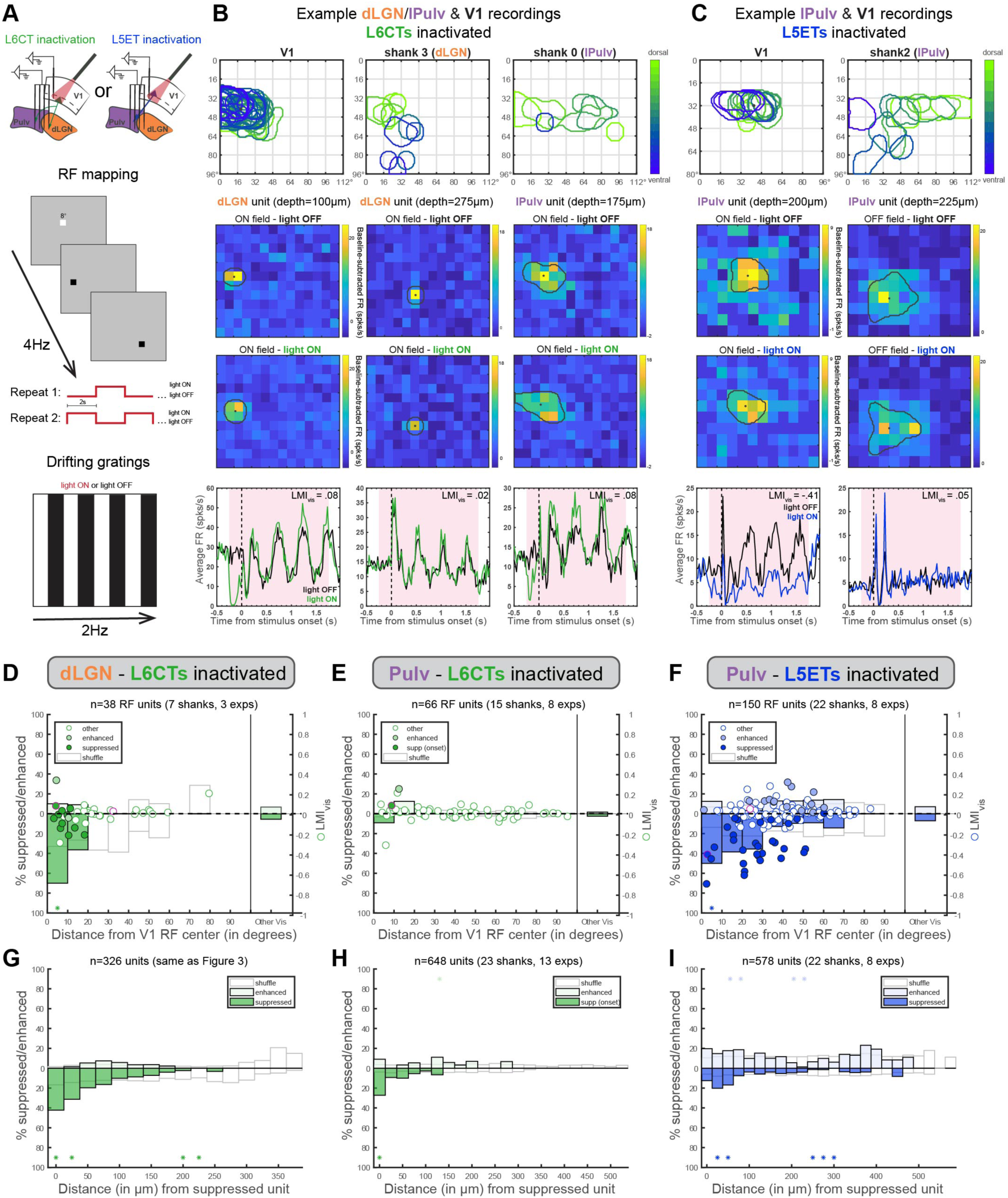
Corticothalamic excitation by both L6CTs and L5ETs is retinotopically organized. (A) Schematics of V1 and thalamus recording configurations (top) and visual stimulation protocols: sparse noise (for receptive field mapping - presented at 4Hz; middle) and drifting gratings (temporal frequency=2Hz) with and without 617nm light for optogenetic CT inactivation (bottom). (B) An example L6CT inactivation experiment in which V1, dLGN and lPulv were recorded from simultaneously. Top: overlayed ON-field RFs (in response to luminance increases) from recording shanks in V1, dLGN and lPulv, colored according to the unit’s relative depth within its respective region. Sparse noise stimuli (8° squares) were presented in a 12×14 grid. Middle: Example dLGN (first two columns) and lPulv (third column) units’ RFs without (top) and with (bottom) L6CTs inactivated, shown as baseline-subtracted FRs at the timepoint of peak response. Overlayed perimeters and dots indicate estimated RF outline and centroids (see Methods). Note that the first dLGN unit’s and the lPulv unit’s RFs overlap with the retinotopic location of V1 recording; the second dLGN unit does not. Bottom: PSTHs of the same units in response to drifting grating stimuli. The retinotopically aligned dLGN and lPulv units’ prestimulus (spontaneous), but not visually evoked, activity was suppressed by L6CT inactivation. (C) Same as (B) but for a L5ET-inactivation experiment in which V1 and pulvinar were recorded from consecutively (10×12 grid for V1 recording, 12×14 for pulvinar). (D) Relationship between light modulation (in drifting grating experiments) and retinotopic displacement from V1 recording site (i.e., the retinotopic locus of L6CT inactivation) from experiments in which sparse noise stimulation was used for both dLGN and V1 recordings. Dots indicate LMI_vis_ (right y-axis) of enhanced, suppressed, and other cells (same as Figure 3 - classified from blank trials). Pink-outlined dots are the example units from (B). Bars indicate proportion of units significantly suppressed or enhanced (left y-axis), binned by retinotopic distance (10° bins). ‘Other Vis’ are all other units from the same experiments whose RFs could not be determined. White bars reflect means of 1000 shuffled distributions (shuffled within experiment). (E) Same as (D) but for pulvinar units defined as “suppressed” from the prestimulus period following light onset (e.g., example pulvinar unit in B). (F) Same as (D) but for pulvinar units during L5ET inactivation experiments. (G) Percent of all unit pairs - consisting of two units, at least one of which was significantly suppressed, recorded on the same shank in the same experiment - in which the second unit was suppressed or enhanced, binned by vertical distance between the channels from which those units were recorded. White bars reflect means of 1000 shuffled distributions (shuffled separately for each recording shank). (H-I) Same as (G) but for pulvinar units in L6CT inactivation experiments (H; “onset suppressed” cells as in E) and pulvinar units in L5ET inactivation experiments (I). Asterisks indicate where actual proportions fell beyond either tail (2.5%) of shuffled distributions.

In both L6CT and L5ET inactivation experiments, the largest suppressive effects (whether on spontaneous and/or visual activity) were observed in thalamic units whose RFs were closely aligned to those recorded in V1, while units whose RF centers were displaced 20° or more were typically unaffected (examples in Figures 4B and 4C; RF distance versus LMI_vis_ for all units with significant RFs in Figures 4D-F). Thus, when considering thalamic units retinotopically aligned (within 10°) to our V1 recordings (and thus confirmed V1 inactivation), at least 50% of dLGN and pulvinar units were classified as “suppressed” by L6CT or L5ET inactivation, respectively, during the drifting grating experiments (Figure 3), compared to 10% and 16% across all visually responsive dLGN and lPulv units (Figures 3D and 3H). This relationship was not observed when we randomly reassigned RF distances among all units recorded within the same experiment (Figures 4D-F, “shuffle” = means of 1000 shuffles). Because of the thalamus’ retinotopic organization, we also found that in the dLGN with L6CT inactivation and pulvinar with L5ET inactivation, units recorded from the same or nearby channels as a suppressed cell were more likely to also be suppressed, and at a significantly greater rate than would be expected from a shuffled distribution (Figures 4G and 4I). This relationship was most striking in the dLGN, where cells’ physical and retinotopic distances are more closely linked than in the pulvinar (Bennett et al., 2019). It is notable that dLGN units whose spontaneous activity was facilitated by L6CT inactivation were typically 50-150µm away from their nearest suppressed cell (Figure 4G), which is consistent with recent reports of L6CTs exerting inhibitory influence over non-retinotopically matched cells (Born et al., 2020). Pulvinar cells whose onset responses (first 200ms) were significantly suppressed by L6CT inactivation (like the example lPulv unit in Figure 4B) also tended to be recorded in closer proximity (Figure 4H). Taken together, these findings show that direct CT excitation, whether from V1 L6CTs or L5ETs to dLGN or pulvinar, is retinotopically organized, even while its influence on visual versus baseline activity depends critically on the CT source (Figure 3).

### Complete L5ET silencing confirms L5 “driving” influence on visual response properties in the pulvinar

Our results thus far demonstrate that V1 L5 and L6 CT projections to the thalamus, while both retinotopic, differ considerably in the extent to which they “drive” or “modulate” visual activity in their thalamic targets. Nevertheless, L5ET inactivation did not completely abolish visual responses in the pulvinar, even in retinotopically aligned cells (e.g., the first unit in Figure 4C, while suppressed by L5ET inactivation, maintained its RF properties). This could mean that V1 L5ETs contribute to but are not wholly necessary for visual responsiveness in the pulvinar. Alternatively, L5ETs in V1 may be necessary drivers but were incompletely inactivated by our optogenetic approach. In fact, we noticed that halorhodopsin was more effective at silencing L5ETs’ spontaneous than visually evoked activity; this could even result in a sharpening, rather than an ablation, of RFs in “inactivated” L5 cells (Figure S4G). We thus sought another optogenetic approach to specifically silence L5ETs with even greater efficacy to more definitively determine the extent to which the pulvinar is driven by V1 L5.

Light-activated chloride channels, such as the blue light-activated, soma-targeted GtACR2 (stGtACR2), have been shown to exhibit enhanced photocurrents relative to other available inhibitory opsins (Mahn et al., 2018). They also have the advantage of providing shunting inhibition as opposed to hyperpolarization (Wiegert et al., 2017) and thereby, potentially, more effective suppression of depolarization caused by visual stimulation. We therefore conducted additional L5ET inactivation experiments in Npr3-Cre mice using stGtACR2 instead of halorhodopsin (Figure 5). While we recorded a similar proportion of inactivated regular-spiking L5 cells in V1 (Figure 5C), those cells were inactivated to a greater degree than when using halorhodopsin. In fact, the majority of these cells were completely silenced (Figure 5D), and this was equally true of their visual responses, including their orientation tuning and RF properties (e.g., Figure 5A), as well as their spontaneous activity (Figure S4I).

**Figure 5.**
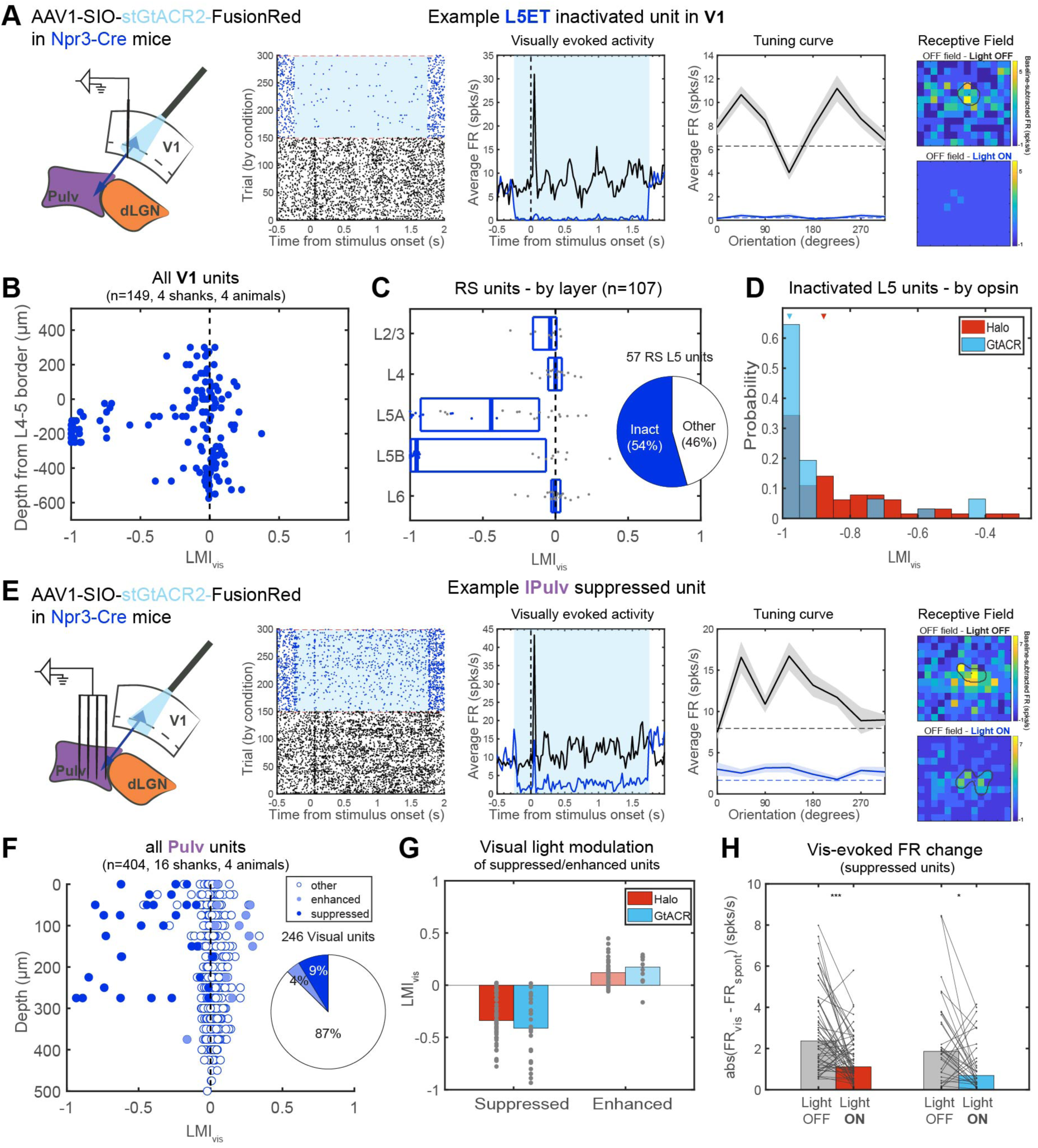
Soma-targeted GtACR2 allows greater L5ET silencing than with halorhodopsin and further demonstrates L5ET “driving” influence over a subset of pulvinar units. (A) stGtACR2-inactivation of L5ETs in V1. Left: schematic of V1 recordings (but were always conducted simultaneously with pulvinar recordings). Right: an example inactivated regular-spiking L5 unit whose spontaneous and visually evoked responses were silenced by blue light. Its orientation tuning and receptive field (RF) were also abolished. Shaded lines in tuning plots depict means and standard errors from each condition, and dotted lines indicate average spontaneous FRs (from blank trials). (B) Light modulation index from visual trials (LMI_vis_) of all recorded V1 units by their depth through cortex, relative to the end of L4. (C) LMI_vis_ of regular-spiking units, by layer. Putative inactivated units are blue dots. Overlay: proportion of all RS L5 units putatively inactivated. (D) Histogram of putatively inactivated units from halorhodopsin (Figure 2) and stGtACR2 experiments. Triangles indicate medians. p=0.0022, Wilcoxon rank-sum test. (E) Pulvinar recordings with L5ET inactivation with stGtACR2. Left: recording and light stimulation configuration. Right: an example pulvinar unit – recorded simultaneously with the example L5ET unit in (A) - whose spontaneous and visually evoked activity, tuning and RF were dramatically suppressed by L5ET inactivation using stGtACR2 (note that its RF aligns with that of the simultaneously recorded V1 unit in A). (F) LMI_vis_ by depth for all recorded pulvinar units (combining lPulv and rmPulv units), classified by effects on their spontaneous activity as significantly suppressed, enhanced or non-modulated (“other”). Overlay: proportion of all visually responsive units in each category. (G). Comparison of suppressed and enhanced units’ LMI_vis_ between halorhodopsin and stGtACR2 inactivation experiments (no significant difference in either suppressed or enhanced cells; p>0.1, Wilcoxon rank-sum tests). (H) Visually induced change in FR in Light OFF versus Light ON (L5ET inactivated) trials in different inactivation experiments (same groups as in G). ***p<0.0001, *p=0.0128, Wilcoxon signed-rank tests.

This pronounced improvement in L5ET inactivation resulted in only marginally greater suppression of a subset of pulvinar neurons (Figures 5G and 5H). Some, such as the example unit in Figure 5E (see also unit #808 in Figure 7C), were deprived of their orientation tuning and RF integrity; this provides definitive evidence of V1 L5ETs “driving” visual response properties in certain retinotopically aligned pulvinar neurons. Still, a similarly small (and even reduced) proportion of all visual pulvinar cells were impacted by complete L5ET silencing with stGtACR2 (Figure 5F) in comparison to silencing with halorhodopsin (Figure 3G). This reduction is likely related to the limited spread of the stGtACR2-encoding AAV, as when considering retinotopically aligned units, the proportion of suppressed cells was about 50%, similar to our halorhodopsin experiments (Figures S4B-D). Therefore, using the potent chloride channel stGtACR2 to silence V1 L5ETs somewhat increased the potency but not the prevalence of suppression in the pulvinar (Figures 5F-H) and more conclusively demonstrates the “driving” influence of L5 CT projections on a subset of pulvinar neurons’ visual activity.

### Combined inactivation of different CT populations shows that individual pulvinar neurons are driven by L5CT, but not L6CT, inputs from V1

Although we have demonstrated different effects of V1 L6CT versus L5ET inactivation on activity in the pulvinar that are consistent with their hypothesized “driving” and “modulatory” roles, the degree of suppression we observed from specific L5ET inactivation – even when L5ETs were essentially completely silenced (Figure 5) - was less complete than in other studies that broadly inactivated all of V1 (Beltramo and Scanziani, 2019; Bennett et al., 2019) or in our own nonspecific cortical inactivation experiments (Figures S5A-F). We thus considered the possibility that while inactivating V1 L6CTs on their own had minimal effect on pulvinar activity (Figure 3F), perhaps they could exert more influence when in concert with the L5ET pathway.

To address this possibility, we took advantage of the fact that AAVretro injected into the pulvinar infects only L5, but not L6, CT neurons (Tervo et al., 2016) (Figure 1B), thus allowing us to express Flp recombinase specifically in corticothalamic L5ET neurons (L5CTs). These injections were made in Ntsr1-Cre mice, where Cre is already present in L6CTs, so that a mixture of AAVs encoding Flp-dependent stGtACR2 and Cre-dependent halorhodopsin injected to V1 resulted in separable opsin expression in each CT population (Figures 6A and 6B). V1 recordings confirmed that blue (455nm) LED light for stGtACR2 specifically silenced units in L5 (Figure 6C), whereas red (617nm) LED light for halorhodopsin inactivated units in L6 (Figure 6D).

**Figure 6.**
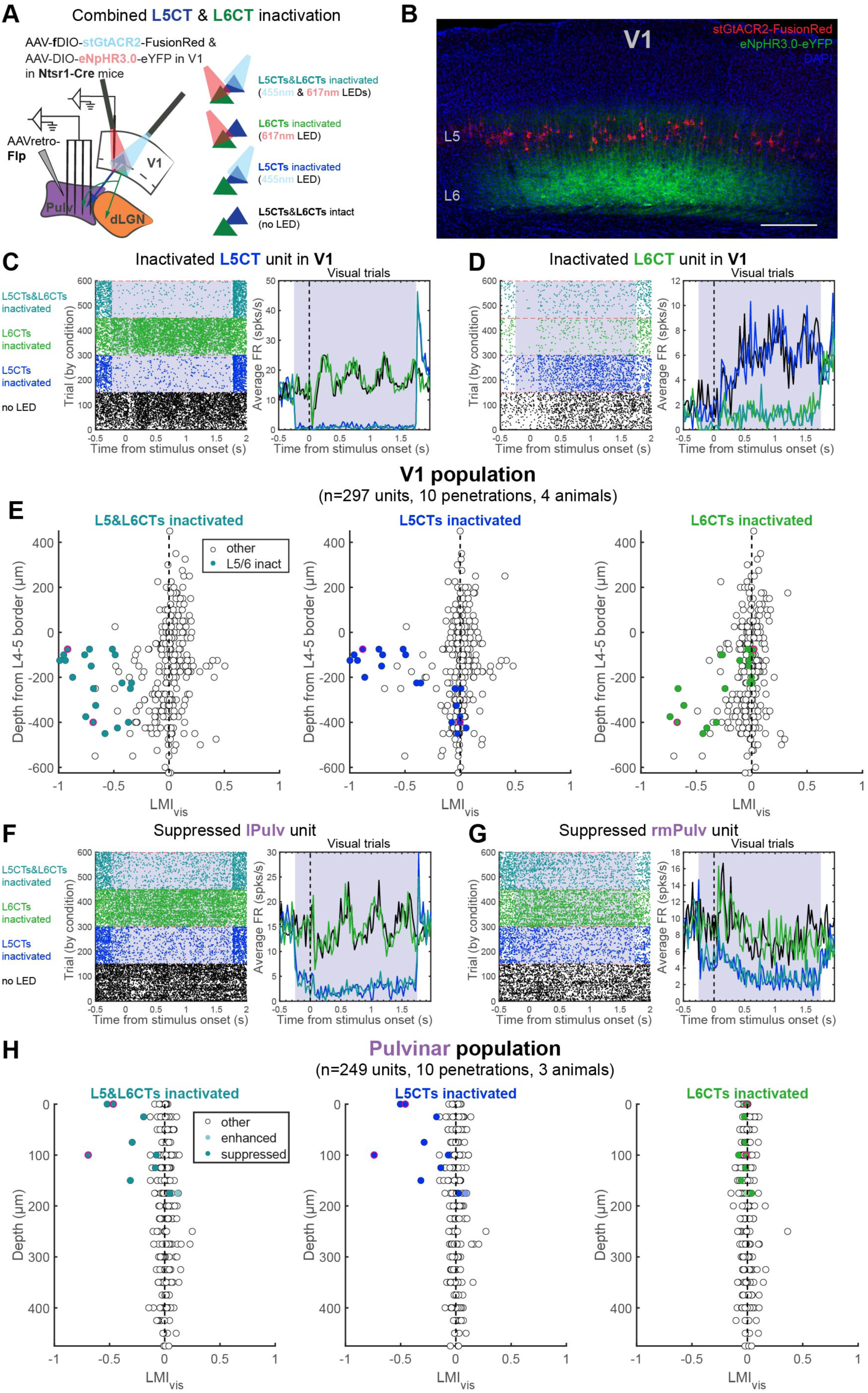
Individual pulvinar neurons are suppressed by V1 L5CT, but not L6CT, inactivation. (A) Experiment schematic. Ntsr1-Cre mice were injected to express Cre-dependent halorhodopsin in L6CTs and Flp-dependent stGtACR2 in L5CTs (L5CT-specific, as opposed to L5ET, expression because of AAVretro-Flp injection into the pulvinar). Right: four different LED stimulation conditions (randomly interspersed). (B) Confocal image (maximum intensity projection) of stGtACR2-expressing L5CTs and eNpHR3.0-expressing L6CTs in V1. Scale bar = 200µm. (C-D) Example L5CT (C) and L6CT (D) inactivated units in V1. Raster plots (left) and PSTHs of average firing rates across visual trials (right). Grey shading indicates the period of LED stimulation. (E) LMI_vis_ calculated from L5CT&L6CT-inactivation (left), L5CT-inactivation (middle), and L6CT-inactivation (right) trials. Colored units are those whose FRs were significantly suppressed by at least 50% in visual trials with combined L5CT&L6CT-inactivation, and dots outlined in pink are the example units in (C) and (D). (F-G) Example lPulv (F) and rmPulv (G) units that were suppressed by L5CT, but not L6CT, inactivation. (H) LMI_vis_ of all pulvinar units across light stimulation conditions of significantly suppressed, enhanced and non-modulated units (classified from blank trials with combined L5CT&L6CT-inactivation). Dots outlined in pink are the example units in (F) and (G).

Fewer L5 units were putatively inactivated in these experiments than when using Npr3-Cre mice, which was expected since opsin expression was restricted to pulvinar-projecting L5ETs (i.e., L5CTs). Across the full depth of V1, combined blue and red LED illumination reduced activity in cells across the infragranular layers, while each LED alone only suppressed units in its corresponding layer (Figure 6E). We confirmed in control animals expressing only one opsin that halorhodopsin was unaffected by our blue LED stimulation, nor was stGtACR2 affected by the red LED (Figures S5G-N).

Using this approach to inactivate L5CTs or L6CTs alone or in combination, we once again observed a subset of pulvinar units that were suppressed by L5CT inactivation (Figures 6F and 6G). Importantly, however, these units were unaffected by L6CT inactivation, and the effect of combined L5CT and L6CT inactivation was indistinguishable from that of L5CT inactivation alone (Figure 6H). While we cannot definitively prove that these same neurons receive input from L6CTs in V1, our prior demonstration of the retinotopic organization of both L5 and L6 CT projections (Figure 4) would support this inference. Thus, these experiments conclusively show that individual pulvinar neurons can be driven by specifically L5, but not L6, CT inputs from V1.

While we have demonstrated that the L5 CT pathway from V1 “drives” a subset of retinotopically aligned pulvinar cells, the full extent of visual activity observed in the pulvinar is not accounted for. When we virally ablated L5ETs (or L6CTs) in V1 to further ensure the complete inactivation of these populations, we still observed robust visual responses in the pulvinar – even in some neurons whose RFs aligned with the cortical area of ablation (Figure S6). Although chronic ablation has the potential to induce compensatory mechanisms, these results are consistent with our optogenetics experiments, where we achieved transient and virtually complete L5ET inactivation and still observed residual visual responses (Figure 5) and even some unaffected yet retinotopically aligned pulvinar cells (Figure S4A). We were therefore interested whether inputs other than from V1 might also play prominent roles in shaping pulvinar activity.

### Cortical and subcortical “driving” pathways converge in the lateral pulvinar

In the lateral pulvinar (lPulv) in particular, an important source of additional input may come from the superior colliculus (SC) (Zhou et al., 2017). Tectopulvinar synapses exhibit morphological and physiological “driver-like” properties with intermediate type 1 “modulator” and type 2 “driver” characteristics (Bickford, 2015; Masterson et al., 2009, 2010). Moreover, recent studies have shown that the SC, but not the cortex, drives activity in the caudomedial pulvinar (cmPulv) (Beltramo and Scanziani, 2019; Bennett et al., 2019). While our experiments with V1 CT manipulations did not concern this region because it does not receive direct V1 input (Zhou et al., 2017), lPulv, which we did record from, is innervated by both V1 and the ipsilateral SC. Although axons emanating from topographically aligned regions of V1 and SC are largely segregated within lPulv (Figure 7A), cortical and SC terminals have been found in close proximity at the electron microscopic level (Masterson et al., 2009; Rovó et al., 2012). Moreover, pulvinar neurons have wide-reaching dendrites that can even stretch across pulvinar subdivisions (Zhou et al., 2017), and short-latency effects of both V1 and SC stimulation on single pulvinar cells have been described (Casanova and Molotchnikoff, 1990). Therefore, we were interested: a) whether the visually responsive cells we recorded in lPulv that were unaffected by L5ET inactivation would be suppressed instead by SC inactivation; and b) whether individual neurons in lPulv might receive convergent L5ET and SC “driving” inputs.

To address these questions, we recorded single-unit activity from lPulv in awake mice expressing stGtACR2 in L5ETs (using Npr3-Cre mice) and halorhodopsin in the SC, thereby allowing us to use different LEDs to inactivate L5ETs and/or SC (Figure 7B). We also made recordings from V1 (data in Figures 5A-D) and the SC (Figures S7E-I) to verify the efficacy of our optogenetic manipulations. Not only were there individual units in lPulv which were strongly suppressed by L5ET inactivation, but we also found units, even in the same recording penetration, which were instead suppressed specifically by SC inactivation (Figure 7C). These example units were “driven” selectively by V1 L5ETs or the SC, as their responses to sparse noise stimuli presented within their RFs were abolished by optogenetic silencing of only their cortical or subcortical inputs, respectively (Figure 7D). Intriguingly, the L5ET-driven unit (#808) was also suppressed by SC-only inactivation, but exclusively in the presence of its preferred direction stimulus (Figure 7C). This suggests that although this unit’s baseline firing and RF is determined by its V1 L5 input, the SC may provide additional input that bestows direction tuning.

**Figure 7.**
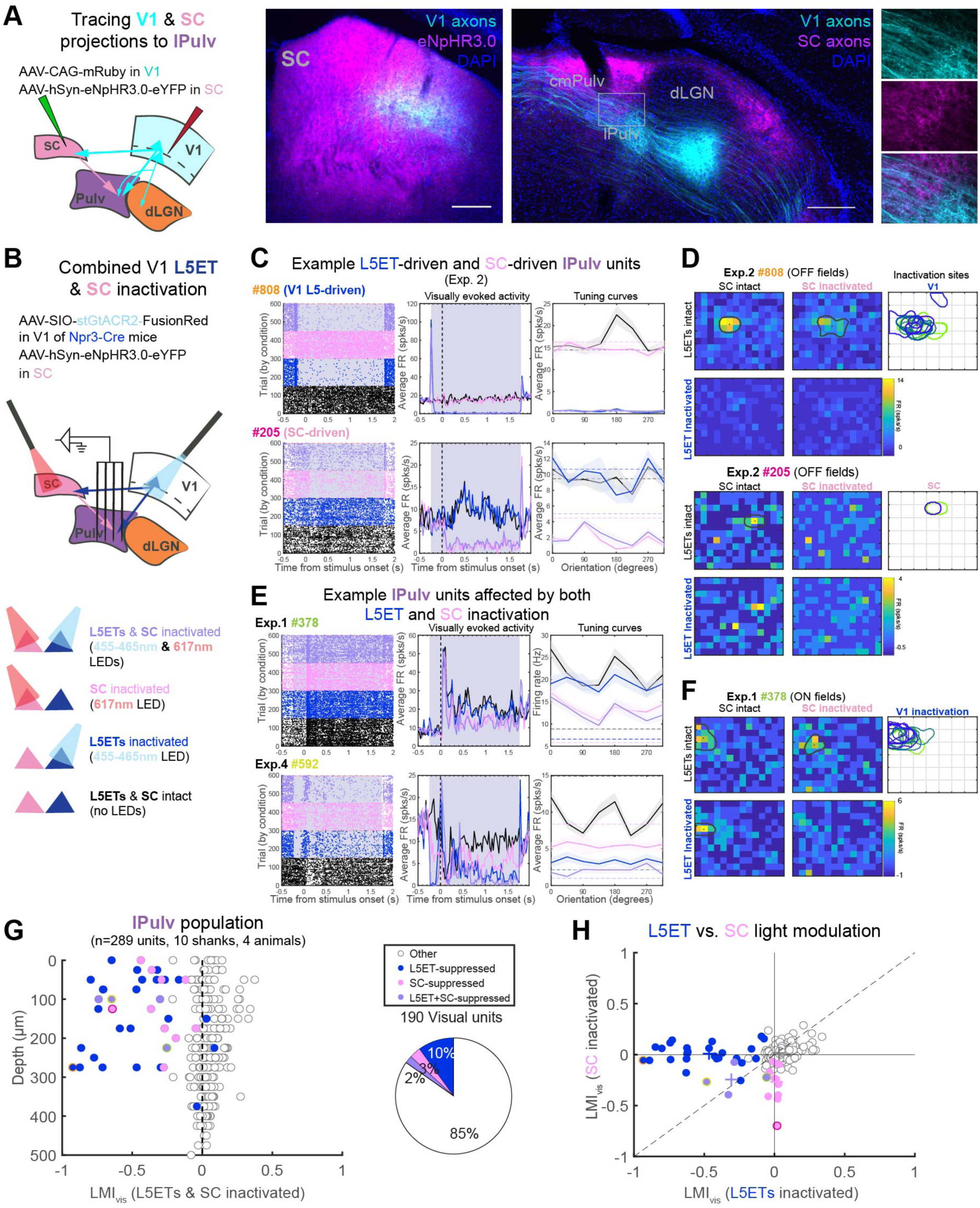
Subsets of neurons in the lateral pulvinar (lPulv) are driven by V1 L5ETs, the superior colliculus (SC), or both. (A) Anterograde tracing of V1 and SC projections to the pulvinar. AAV injection of eNpHR3.0-eYFP overlaps with V1 axons in the SC (middle), yet cortical and SC axons are largely segregated with minor overlap in lPulv (right, with higher magnification of boxed area in each channel at far right). Scale bars = 200µm. (B). Experimental schematic for dual-optogenetic inactivation of V1 L5ETs and SC during pulvinar recordings. Bottom: four inactivation conditions. (C) Two example lPulv units, recorded on the same shank in the same experiment. From left to right: raster plots with trials organized by inactivation condition; PSTHs of average visually evoked FRs; and average FRs in response to different drifting grating orientations (shaded lines depict means and standard errors, dashed lines indicate spontaneous FRs). The first unit (#808) was suppressed by L5ET inactivation conditions, whereas #205 was suppressed by SC inactivation conditions. (D) Receptive fields (RFs) of units from (C) under different inactivation conditions. #808’s RF was abolished by L5ET but not SC inactivation, while the opposite was true of #205. RF maps depict average baseline-subtracted firing rates at the timepoint of peak response across all OFF stimulus (black square) positions. Outlines depict estimated RF boundaries (see Methods) under each condition. Right: overlayed RFs of V1 (upper) and SC (lower) units recorded in the same animal. (E) Example units that exhibit combined effects of L5ET and SC inactivation. Same plot descriptions as for (C). (F) Unit #378’s (from E) RF under different inactivation conditions. Right: Overlayed RFs of simultaneously recorded V1 units. SC was not recorded from in this animal. No RF information from Exp. #4 for unit #592. (G) LMI_vis_ from combined L5ET & SC inactivation trials, by depth. Units are classified according to whether their visual and/or spontaneous activity was significantly suppressed by SC inactivation, L5ET inactivation, or both (see Methods for details). Color-outlined dots correspond to example units in (C) and (E). Right: proportion of different suppressed unit classes out of all visually responsive units. (H) LMI_vis_ for each unit in L5ET inactivation versus SC inactivation conditions (same unit classification as in G). Crosses indicate medians of each unit class.

Across our recorded population, we observed a wide range of effects of V1 L5ET versus SC silencing (Figure 7G). Despite the dorsal-ventral distribution of SC and V1 axons in lPulv (Figure 7A), units whose visually evoked and/or spontaneous firing rates were significantly suppressed only by L5ET inactivation or by SC inactivation (L5ET-suppressed or SC-suppressed, respectively) were fairly dispersed (Figure 7G). While most of the modulated units were only L5ET- or SC-suppressed (i.e., units falling along either axis in Figure 7H), a handful of units were suppressed under both L5ET and SC inactivation conditions (lower left quadrant of Figure 7H).

Indeed, we found a number of intriguing instances of combinatorial effects of L5ET and SC silencing on individual units, in addition to the direction-tuned cell in Figure 7C (#808). The first example unit in Figure 7E was mainly impacted by SC, but not L5ET, silencing in its response to drifting grating stimuli, yet its RF was influenced by both SC and V1 L5 inputs (Figure 7F). Similarly, the second unit in Figure 7E exhibited suppressed spontaneous and visually evoked responses from either SC or L5ET inactivation and was even more suppressed by their combined inactivation, suggesting some degree of visual “drive” coming from both cortical and subcortical sources. We made similar observations of cortical and subcortical inactivation having varied and sometimes combinatorial effects on individual units in additional experiments in which V1 (rather than L5ETs specifically) was inactivated (Figures S7A-D).

We have therefore shown that V1 L5ETs are not the only excitatory projections to “drive” visual activity in this region of the HO visual thalamus. Indeed, other units are driven by SC projections, and some even appear to receive input from both sources that convey varying degrees of visual drive. Given our previous demonstration of the retinotopic specificity of L5ET excitation (Figure 4), we may well underestimate the extent of combined cortical and subcortical influences (e.g., in Figure 7D, unit #205’s RF aligned with the SC but not L5ET inactivation centers). Regardless, we have shown that while V1 CT projections from L5 – but not L6 – provide retinotopic, visual driving input to the pulvinar, the SC can also provide some degree of visual drive. These cortical L5 and subcortical inputs thereby converge within the lateral pulvinar - and even onto individual neurons – to shape visual responses.

## Discussion

What roles do two, seemingly parallel corticothalamic pathways originating from different layers of the primary visual cortex play in thalamic processing in an awake animal? The distinct characteristics of L5 versus L6 CT inputs have for decades led to the idea that these are not redundant pathways but rather distinct “driving” versus “modulatory” projections (Rouiller and Welker, 2000; Sherman and Guillery, 1996, 1998). While this idea has been highly influential to the study of corticothalamic circuit organization and function, it has never been directly tested. This is largely due to technical limitations in selectively silencing one or the other CT pathway and assessing downstream effects on the thalamus in an awake animal. Taking advantage of cell type-specific transgenic mouse lines, optogenetics and high-density, multielectrode recordings of single-unit activity in awake mice, we find pronounced differences in how these populations influence sensory responses in the visual thalamus. While both L5 and L6 CT projections from V1 provide retinotopic excitation onto their thalamic targets, the extent to which that excitation imparts visual information differs considerably. Our inactivation studies demonstrate that V1 L6CTs can provide some degree of baseline drive, mainly onto retinotopically aligned cells of the dLGN, yet are not required for visual responses in either FO or HO visual thalamus. In contrast, silencing the L5CT pathway from V1 suppressed visually evoked as well as baseline activity in many retinotopically matched pulvinar neurons, some of which relied entirely on that input for their tuning and/or RF properties (e.g., Figures 5E and 7C-D). Therefore, our results affirm a longstanding hypothesis that the L5, but not the L6, CT pathway from V1 constitutes a functionally “driving” pathway that conveys visual information to the rodent pulvinar, whereas the L6 CT pathway is fundamentally “modulatory”.

While we were able to selectively inactivate L5ET versus L6CT subpopulations by using different transgenic mouse lines, there were still some limitations to the specificity of our approach. For instance, although the Ntsr1-Cre line is selective for L6CTs that project to the dLGN (Bortone et al., 2014), not all project to the pulvinar (Figure 1E). If anything, this might have led us to overestimate the influence of V1 L6CTs in the pulvinar, yet we still saw minimal impact of their inactivation. On the other hand, many L6 projections to the rodent pulvinar come from extrastriate areas (Blot et al., 2020; Roth et al., 2016; Scholl et al., 2020; Souza et al., 2020) (Figure 1B), which were not targeted by our injections. Our findings of fundamentally similar “modulatory” influences of V1 L6CTs in the dLGN and pulvinar (from our present inactivation and prior activation studies; Kirchgessner et al., 2020) would lead us to expect that the cumulative effect of L6CT inactivation across cortical areas on the pulvinar would be similar to what we observed in this study in the dLGN (which gets the majority of its L6CT input from V1; Bienkowski et al., 2019; Briggs, 2020) – namely, suppressed spontaneous but not visual activity. Since L6CTs themselves have low spontaneous firing rates (Figure S2C), we attribute their effect on baseline thalamic activity to the high degree of convergence of many L6CTs onto single thalamic cells (Reichova and Sherman, 2004; Sherman, 2016) and to the fact that our recordings were conducted in awake animals as opposed to under anesthesia, which reduces spontaneous firing rates in the thalamus (Durand et al., 2016). Additionally, our approach most likely excluded L6b cells that exclusively project to HO thalamus and exhibit type 2 (driver), rather than type 1 (modulator), characteristics (Ansorge et al., 2020; Hoerder-Suabedissen et al., 2018). Although we did not observe many pulvinar-projecting L6CT neurons that might fall within this category (retrogradely labeled tdT-negative cells in Ntsr1-Cre/Ai14 mice, Figure 1E), whether they would more closely resemble L5 “drivers” or other L6 “modulators” in terms of their contributions to functional response properties in the pulvinar in vivo is an open question.

Meanwhile, our primary strategy for silencing L5 CT projections was to use the Npr3-Cre mouse line to restrict inhibitory opsin expression to L5ET neurons in V1. This offers a marked improvement in specificity from other studies that have relied on non-specific cortical inactivation (Beltramo and Scanziani, 2019; Bender, 1983; Bennett et al., 2019; Casanova et al., 1997), which would be expected to also suppress activity in other cortical areas (Leopold, 2012) and thus confound the driving influence of V1, specifically L5ETs, on the pulvinar with that of extrastriate cortex. Indeed, we observed somewhat more widespread pulvinar inactivation when we non-specifically inactivated cortex, even when that inactivation was confined to V1 through AAV injection (FigureS S5A-F). Still, not all of our inactivated L5ETs project to the pulvinar (Figure 1F), as these neurons can innervate any combination of multiple subcortical areas (including pons and SC, Figure S1, and striatum (Kim et al., 2015)). In fact, in light of ours (Figure 7) and others’ demonstrations of SC driving influence on the pulvinar (Beltramo and Scanziani, 2019; Bennett et al., 2019), it is possible that some of the suppression we observed in the pulvinar from L5ET inactivation could be mediated indirectly by the SC. However, we still observed suppressed pulvinar activity when we restricted our L5 inactivation to L5CTs (Figure 5) and found largely separable effects of SC and L5ET inactivation in the lateral pulvinar (Figures 7 and S7A-D) and even within the SC itself (Figures S7E-I). Altogether, this would suggest that L5ETs do not rely on an indirect pathway through the SC to drive visual responses in the pulvinar. Another caveat to our L5ET and L5CT silencing approaches is that our area of inactivation was limited by the spread of our AAVs (Figure S4B) and likely did not encompass all of V1. However, this limitation was addressed by assessing the effects of L5ET inactivation on retinotopically aligned pulvinar neurons (Figures 4 and S4C-D). The fact that half of those cells were still unmodulated by V1 L5ET inactivation suggests that there may be other “driving” inputs to the pulvinar.

Indeed, the pulvinar is innervated by a number of other cortical and subcortical inputs whose in vivo functional influences are not yet known. First, it receives inhibitory input from areas like the ventral LGN (vLGN), the zona incerta and the anterior pretectal nucleus – all of which receive collaterals from L5ETs (Barthó et al., 2002; Blot et al., 2020; Bokor et al., 2005; Bourassa and Deschênes, 1995; Halassa and Acsády, 2016). These inhibitory pathways might underlie some of the excitatory effects we observed in the pulvinar with L5ET inactivation (Figure 3H). Meanwhile, the rodent pulvinar also receives excitatory L5 input not just from V1, but also from extrastriate areas (Blot et al., 2020; Roth et al., 2016; Scholl et al., 2020; Souza et al., 2020) (Figure 1B). Based on our observations of some retinotopically aligned pulvinar units that were unaffected by L5ET inactivation (Figures 4F and S4A) or ablation (Figure S6) – similar to what was observed in the cat following V1 cooling (Casanova et al., 1997) - we predict that L5ETs in extrastriate areas may provide additional driving inputs. Further work is needed to conclusively address this possibility and whether L5 inputs from multiple cortical areas could even provide combined driving input to some neurons.

Our results also show that subcortical excitatory inputs from the SC can intermingle with L5 driving inputs within the lateral pulvinar and even converge onto individual neurons. In fact, while relatively rare, we found cases of units whose firing rates, tuning and/or receptive fields were influenced by both L5ET and SC inactivation (Figures 7 and S7A-D). This would suggest that some pulvinar neurons integrate information conveyed by cortical (L5) and subcortical (SC) sources. This inference is supported by prior reports of cortical and subcortical stimulation having combined effects in the cat LP-pulvinar complex (Benedek et al., 1983) and in the rodent HO somatosensory thalamic nucleus, POm (Groh et al., 2014). Given the role of the SC in processing object motion, eye movements, and spatial attention (Krauzlis et al., 2013), integration of various visual, motor, and other signals within the pulvinar could play a key role in sensory-guided behaviors. This idea is in line with evidence that the rodent pulvinar conveys contextual information (e.g., a mismatch between visual flow and running speed) to extrastriate areas (Blot et al., 2020) and even back to V1 (Roth et al., 2016). It is notable, however, that SC inactivation/lesion was shown to considerably impair visual responses in regions of the pulvinar homologous to the mouse lPulv (Baldwin et al., 2017) in rabbits (Casanova and Molotchnikoff, 1990), but not in macaques (Bender, 1983). It is therefore possible that subcortical inputs might play a more prominent role in shaping pulvinar visual response properties in non-primates (Casanova and Molotchnikoff, 1990).

Altogether, our results demonstrate a functional dissociation between L6 and L5 corticothalamic pathways that has long been supposed but never directly proven. Many notable distinctions between FO and HO thalamic nuclei – such as their excitatory input sources (Sherman and Guillery, 1996), relative proportions of different synapse types (Horn and Sherman, 2007; Horn et al., 2000; Wang et al., 2002), transcriptional profiles (Phillips et al., 2019), and functional properties (Halassa and Sherman, 2019) – could have led “driver” and “modulator”-type inputs to differently contribute to activity in the HO compared to FO thalamus. Instead, we find that the effects of inactivating V1 L5 versus L6 CT pathways on visual responses in the pulvinar of awake mice are consistent with their hypothesized “driving” versus “modulatory” functions. Given the similarities in CT synapse types (Rouiller and Welker, 2000) and the consistent effects of cortical lesion/inactivation on sensory activity in the HO thalamus across systems and species (Beltramo and Scanziani, 2019; Bender, 1983; Bennett et al., 2019; Casanova et al., 1997; Diamond et al., 1992; Mease et al., 2016), we suggest that this functional distinction between L5 and L6 CT projections to HO thalamus is a common organizational principle of corticothalamic function. We also found that the superior colliculus can play important “driving” roles in the rodent pulvinar and can even converge with the L5 cortical driving pathway; the extent to which this, too, generalizes across species will be of considerable interest for future studies of pulvinar function. Our results thus illuminate how sensory information is routed to the HO thalamus through distinct cortical (and subcortical) channels, which has important implications for understanding how the HO thalamus supports sensation and behavior in the awake animal.

## Acknowledgements

We thank Dr. Hongkui Zeng (Allen Institute) for informing us about and sharing the Npr3-IRES-Cre-neo line used in this study, as well as Dr. Farran Briggs and Dr. Sonja Hofer for providing feedback on the manuscript. We also thank other members of the Callaway Lab, especially Dr. Celine Vuong for making the fDIO-stGtACR2 plasmid and Rachel Cassidy for comments on the manuscript, as well as the Salk Institute’s Viral Vector and Advanced Biophotonics Cores for additional support. This work was supported by NIH grants F31 EY028853 (M.A.K.), R01 MH063912 and R01 EY022577 (E.M.C.).

## Author contributions

M.A.K. and E.M.C. conceptualized and designed the experiments. M.A.K. collected and analyzed the data. L.D.F. provided technical assistance with histology, immunohistochemistry and injections surgeries and contributed to data processing and analysis (spike-sorting and cell counting). M.A.K. and E.M.C. wrote the manuscript.

## Methods

### Resource availability

#### Lead Contact

Further information and requests for resources and reagents should be directed to and will be fulfilled by the Lead Contact, Edward Callaway.

#### Materials availability

Plasmid for Flp-dependent stGtACR2-FusionRed (pAAV-Ef1a-fDIO-stGtACR2-FusionRed) that was generated for this study will be made available upon request to the Lead Contact.

#### Data and code availability

The code generated for this manuscript will be made available on GitHub (https://github.com/SNLC). The data will also be made available.

### Animals

The following transgenic mouse lines were used in this study:

- Ntsr1-Cre GN220 (Gong et al., 2007) - L6CT inactivation and combined L5CT & L6CT inactivation experiments
- Npr3-IRES-Cre-neo (Daigle et al., 2018) - L5ET-inactivation experiments
- PV-Cre and Dlx5-Flp/CCK-IRES-Cre - V1 inactivation by activating PV+/Dlx5+ inhibitory interneurons
- PV-Cre/Ai32 - unrestricted V1 inactivation by activating constitutively ChR2-expressing PV interneurons
- Ntsr1-Cre/Ai14 - L6CT quantification

Cre-negative animals were also used for control physiology experiments and anatomy experiments not requiring Cre-dependent AAV expression (e.g., Figures S1F-H). Both female and male mice were used between 8-17 weeks of age (more commonly 9-14 weeks) at the time of AAV injection and <20 weeks at the time of in vivo recordings. All experimental procedures followed protocols approved by the Salk Institute Animal Care and Use Committee.

### Methods

#### Surgeries

For injections, mice were anesthetized with a ketamine/xylazine cocktail (100mg/kg ketamine, 10mg/kg xylazine) via intraperitoneal injection and secured in a stereotax (David Kopf Instruments Model 940 series). A small craniotomy was made at the injection site, and a glass pipette (24-30µm tip diameter) was lowered to the desired depth and was left for 3-5 minutes before injecting. AAV or a cholera toxin subunit B-Alexa Fluor conjugate (see below) was pressure-injected with a syringe at an approximate rate of 20µl/minute and let rest for at least 5 minutes (10+ minutes for subcortical injections) before raising to prevent backflow. The following stereotactic coordinates were used for injection sites (all left hemisphere, in mm relative to bregma; A/P, M/L, D/V):

- V1: -3.2, -2.65, -0.64-0.3 (2 depths)
- lPulv: -1.6, -1.8, -2.45
- rmPulv: -1.05, -1.2, -2.55
- SC, medial site: -3.5, -0.45, -1.1-1.2; SC, lateral: -3.5, -0.8, -1.25-1.3
- Pons: -3.5, -0.85, -5.75-5.9
- lateral visual cortex (VisL): -2.6-2.75, -3.8, -0.45-0.3 (2 depths)

At the end of injections, the skin incision was closed with Vetbond (3M) and mice were given a subcutaneous injection of buprenex (0.5-1.0 mg/kg) and Ibuprofen in their water bottles.

At least four days and up to 1 week prior to recordings, mice underwent an acute surgery for headframe implantation. Under isoflurane anesthesia, skin was cut away and a circular headframe (7-mm inner diameter) was secured to the skull with dental cement (C&B-Metabond, Parkell). A dull pipette attached to a micromanipulator (MP-285, Sutter Instrument) was used to relocate bregma and mark positions with a waterproof pen for aiming craniotomies for thalamus recordings (coordinates relative to bregma: 1.50-2.75mm lateral, 1.8-1.9mm posterior for lPulv, 1.0-2.0mm lateral, 1.0mm posterior for rmPulv). The exposed skull was covered with a silicone elastomer (Kwik-Cast, World Precision Instruments) and mice were given a subcutaneous injection of buprenex (0.5-1.0 mg/kg), Ibuprofen in their water bottles, and at least one full day undisturbed in their cages.

#### Viruses

The following AAV vectors were used for optogenetics experiments in this study:

- AAV5-EF1a-DIO-eNpHR3.0-eYFP (120-200nl, 3-4e12 GC/ml, UNC) - for L6CT/L5ET halorhodopsin inactivation
- AAV1-hsyn1-SIO-stGtACR2-FusionRed (120-200nl, 0.9-1.8e12 GC/ml, Addgene) - for stGtACR2 inactivation
- AAV5-Ef1a-fDIO-stGtACR2-FusionRed (200-240nl, 1.77e12 GC/ml, Salk Vector Core) - for L5CT inactivation
- AAV5-hsyn-eNpHR3.0-eYFP-WPRE-pA (60-70nl, 5.5e12 GC/ml, UNC) - for SC inactivation
- AAV1-EF1a-fDIO-hChR2(H134R)-EYFP (150nl, 7.78e11 GC/ml, Salk Vector Core) - V1 inactivation in Dlx5-Flp/CCK-IRES-Cre mice
- AAV1-EF1a-DIO-hChR2(H134R)-eYFP.WPRE.hGH(Addgene20298P) (100nl, 8.4e12 GC/ml, UPenn Vector Core) - V1 inactivation in PV-Cre mice

The Cre-dependent halorhodopsin AAV was injected into V1 of Ntsr1-Cre mice and was allowed up to 6 weeks for expression (as well as for non-specific halorhodopsin expression in the SC). However, slightly smaller (120-150nl) halorhodopsin injections were made in Npr3-IRES-Cre-neo mice and restricted to 3 weeks expression time because expression in other layers and an unhealthy-looking L5 appeared at longer timepoints and/or with larger injection volumes in this mouse line. Consequently, animals in which axons were visible in areas not targeted by L5ETs (e.g., TRN) were excluded.

Viruses for anterograde tracing were:

- AAV5-CAG-FLEX-GFP (120-160nl, 5.2GC/ml, Salk Vector Core)
- AAV5-EF1a-DIO-eYFP (120-180nl, 2-4e12 GC/ml, UNC)
- AAV8-CAG-mRuby2 (100nl, 2.4 GC/ml, Salk Vector Core)

For Flp-dependent stGtACR2 expression in L5CTs, we injected 20-40nl of AAVretro-EF1a-Flp (3.6e11, Salk Vector Core) into both lPulv and rmPulv injection sites. For retrograde tracing of L6CTs, we injected 40nl of cholera toxin subunit B (CTB) conjugated to Alexa Fluor 488 (0.5%) or 647 (0.25%; Life Technologies) into the pulvinar. In 2/3 of these experiments, we injected one of each of these CTBs into lPulv and rmPulv sites (remaining animal was only injected in lPulv). For retrograde tracing of L5CTs, we used a self-complimenting (sc)AAVRetro-hSyn-mCherry (1.51e12 GC/ml, Salk Vector Core) because we observed more widespread and brighter labeling of L5CTs than with CTB. 20nl were injected into both lPulv and rmPulv injection sites.

For diphtheria toxin experiments, we injected 120-170nl of a mixture of AAV8-mCherry-flex-dtA (3.7 GC/ml, UNC) and either of the DIO-eYFP/GFP anterograde viruses (above, at 1:1 ratio; volume split across two V1 injection sites in one Npr3-Cre animal), as well as a 1:2 dilution of the same anterograde virus at the same injection sites and volumes in the right hemisphere.

Mice were recorded from 10-15 days later.

#### In vivo electrophysiology

Following 2-4 days of habituation to the running wheel at the recording rig (30+minute sessions), mice were anesthetized with isoflurane to make craniotomies above the desired recording sites. Mice were then head-fixed on their wheel, where they were free to run at their will and their movement was tracked with a rotary encoder. A black curtain was secured around the headframe holder to prevent the mice from seeing light from optogenetic stimulation. Individual recording sessions typically lasted 2-3 hours. In the case of multiple recording sessions from a single animal, mice were given at least 30 minutes and up to a day undisturbed in their cages with food and water between sessions.

For our recordings, we used 64- or 128-channel silicon microprobes (Du et al., 2011; Yang et al., 2020) connected to an Intan RHD2164 128-channel amplifier board. Data was acquired at 20kHz with the OpenEphys data acquisition system (Siegle et al., 2017). For the majority of recordings, V1/SC and thalamus recordings were conducted simultaneously using two separate 128-channel amplifier boards (thus 192-256 channels recorded simultaneously through OpenEphys). For V1 and SC recordings, we used either 64D (64 channels on one shank, 1.05mm vertical extent of electrodes) or 128AN (two shanks each with 64D configuration, separated by 300µm) probe configurations. For thalamus recordings, we always used a 4-shank probe (128DN or 128D; 32 channels per shank over 775μm depth, 150μm or 330μm separation between shanks, respectively). All probes were gold-electroplated with an Intan RHD electroplating board to reach ∼0.2MΩ electrode impedances.

Probes were lowered slowly into the brain with a digital and/or manual micromanipulator (MP-285, Sutter Instrument Co; 1760-61, David Kopf Instruments also used for simultaneous V1/SC recordings, 80° orientation) to approximate depths of 1.1mm (V1), 1.5mm (SC), or 2.5mm (pulvinar). Agarose (∼3.5%; A9793, Sigma-Aldrich) was then poured to fill the well of the headframe holder, thus covering the probe shank(s) and the tip of the optical fiber(s); this helped with grounding as well as recording stability. After lowering, the probes were left in place untouched for at least 30 minutes before data acquisition commenced. For all recordings, probes were coated with a 1-2.5% solution DiI or DiD (D282 or D7757, Thermo Fisher) in order to verify recording locations post-mortem. For thalamus recordings, probes were typically lowered once without DiI/DiD to check their location (based on presence or lack of highly temporally modulated visual responses, which indicated placement in dLGN) before being raised and relowered with DiI/DiD (for some experiments using 128D probes, they were not raised and relowered with DiI/DiD in the same position until after data acquisition was complete).

#### Visual stimulation

Visual stimuli were generated through custom MATLAB code using Psychtoolbox and presented on a 24” LED monitor (GL2450-B, BenQ). The monitor screen was positioned 12cm from the mouse’s right eye. For drifting gratings experiments, full-field square-wave gratings were presented at four orientations in eight directions, 0.04 cycles/° spatial frequency, and 2Hz temporal frequency. A full trial consisted of a 0.5-second pre-stimulus period (grey screen), 2 seconds of drifting grating presentation, and 1.5-2 seconds post-stimulus period (grey screen). 20% of trials were “blank” trials, in which the screen remained grey for the full trial duration; these trials were used for assessing effects on spontaneous activity (see Analysis section below). Each unique drifting grating stimulus was presented at least 16 times under each light stimulation condition across the full experiment.

For receptive field mapping, a sparse noise stimulus protocol was used in which individual black or white 8° squares were displayed at random on a grey background anywhere within a chosen areal grid (10×12, 12×14 or 14×16). Each square was presented for 250ms. A “trial” consisted of both black and white squares being presented exactly once at all possible stimulus locations, and the same pattern of stimulus presentations was repeated in each light stimulation condition (see below). Timings of individual stimulus presentations were identified with a photodiode attached to the upper left corner of the monitor, whose luminance changed at each new stimulus presentation (i.e., every 250ms). Across the full experiment, each stimulus (black or white square) was presented at each location in the grid 12-24 times under each light stimulation condition.

For V1 recordings, an additional 2-minute run of 2-second full-field screen flashes was presented at the beginning and end of the recording session. These were used for current source density analysis to identify cortical layers (see Analysis section below).

#### Optogenetic stimulation

Optogenetic stimulation was controlled via an Arduino Zero microcontroller board, which interfaced with MATLAB through a serial port connection. A 12-bit DAC and/or a digital output pin was connected to an LED driver (ThorLabs T-Cube LED driver or Plexon PlexBright LD-1 single channel LED driver) or a laser module (635nm collimated diode laser, Laserglow Technologies). LEDs were used for the majority of experiments, but the 635nm laser was used for five L6CT-halorhodopsin experiments. For other halorhodopsin stimulation experiments, we used a 617nm fiber-coupled LED module (ThorLabs), and for stGtACR2 and ChR2 experiments we used a 465nm (PlexBright, Plexon) or a 470nm (ThorLabs) fiber-coupled LED module. Alternatively, a 455nm LED (ThorLabs) was used for combined halorhodopsin and stGtACR2 CT inactivation experiments (Figure 5) to avoid the blue LED falling within halorhodopsin’s excitation range. LED/laser light was outputted through a custom optical fiber patch cord (1mm diameter, 0.39 NA, ThorLabs; or 400µm diameter, 0.39NA, ThorLabs for dual-optogenetics experiments) and positioned as close as possible to the pial surface without hitting the recording probes. Light powers measured from the optical fiber tip (with PM100D with S121C power sensor, ThorLabs) were: 4-5.5mW from 1mm fiber or 0.75-1.0mW from 400µm fiber for halorhodopsin stimulation (617nm LED or 635nm laser); 1mW from 1mm fiber or 0.12-.16mW from 400µm fiber for L5-stGtACR2 stimulation (455nm or 465nm LEDs); 2.2mW from 1mm fiber or 0.25mW from 400µm fiber for L6CT-stGtACR2 stimulation (455nm or 465nm LEDs); 5-10mW from 1mm fiber for AAV-targeted ChR2-stimulation of interneurons for V1 inactivation (470nm LED); and 0.8-1.1mW from 1mm fiber or 0.46mW from 400µm fiber for V1 inactivation in PV-Cre/Ai32 mice (465nm or 470nm LEDs).

In drifting grating experiments, light stimulation began 250ms after the trial start and 250ms prior to the visual/blank stimulus presentation and stayed on for 2 seconds. All drifting grating stimuli (including “blank” stimuli) were presented an equal number of times under every light stimulation condition (including no light). Trials in different light stimulation conditions were randomly interleaved throughout the experiment.

In receptive field mapping experiments, light alternated between ON and OFF in 2 second intervals for the full trial duration, and each unique pattern of sparse noise stimuli was presented with all possible patterns of light stimulation (e.g., 1^st^ repeat: light ON, light OFF…; 2^nd^ repeat: light OFF, light ON…). For dual-optogenetic experiments, there were four possible light stimulation patterns: 1) LED#1 ON and LED#2 OFF, LED#1 OFF and LED#2 ON…; 2) LED#1 OFF and LED#2 ON, LED#1 ON and LED#2 OFF…; 3) LEDs#1&2 ON, LEDs#1&2 OFF…; 4) LEDs#1&2 OFF, LEDs#1&2 ON….

#### Histology

At the end of recordings, animals were given an intraperitoneal injection of euthasol (15.6mg/ml) and perfused with phosphate-buffered saline (PBS) followed by 4% paraformaldehyde. Brains were dissected out, post-fixed in 2% PFA and 15% sucrose solution at 4°C for ∼24 hours, moved to 30% sucrose at 4°C for another ∼24 hours, and then frozen in sucrose and sliced on a freezing microtome. Brains were sliced coronally into 50μm sections, starting from the anterior edge of the hippocampus to the posterior end of cortex. All sections were counterstained with 10μM DAPI in PBS before being mounted and cover-slipped with Polyvinyl alcohol mounting medium containing DABCO. Additional immunohistochemistry for calretinin was performed on thalamic sections (except in cases with DiD and both red- and green-fluorescent axons in the thalamus; e.g., combined SC and L5/V1 inactivation experiments) by incubating at 4°C for 16-20 hours with rabbit anti-calretinin primary antibody (1:1000; Swant 7697) in 1% Donkey Serum/.1% Triton-X 100/PBS, followed by donkey anti-rabbit conjugated to Alexa 647 (1:500; A-31573, Life Technologies) before DAPI counterstaining. Imaging was performed on an Olympus BX63 microscope (10x objective for most images, 20x for cortical z-stacks for cell counting) or a Zeiss LSM880 confocal microscope (20x).

### Quantification and statistical analysis

#### Data Processing

We used Kilosort2 (Pachitariu et al., 2016) to semi-automatically spike-sort extracellularly recorded data. First, different recordings from the same recording session (drifting gratings and RF mapping experiments) were concatenated together into a single binary file. We then subtracted out the large voltage deflections caused by light onsets and offsets prior to high-pass filtering (>150Hz) in Kilosort2 because otherwise we found that spike detection within ∼10ms of the light onset/offset times was compromised. Additional “spikes” were removed within 1.5ms of the light onset/offset times to ensure that no further optogenetic artifacts were included in the analyses.

Phy2 was used to validate and further curate as needed the clusters outputted by Kilosort2. In some experiments where optogenetic artifacts couldn’t be completely removed in Kilosort2, they were typically easily removed in Phy2 because they appeared as outliers in its principal component features view. Units were considered “good” single-units whose waveforms clearly deviated from the noise, had a clear refractory period in their auto-correlogram, and no evident refractory period in their cross-correlogram with other units. From there, “good” units were included in subsequent analyses which had fewer than 0.5% refractory period violations, visually evoked and/or spontaneous firing rates >=0.25Hz, and “unit quality” (isolation distance (Schmitzer-Torbert et al., 2005)) greater than 16.

#### Unit classification

For thalamus recordings, the fluorescent traces left by the lipophilic dyes (DiI or DiD) on the probe shanks were identified in histology and used to assign all units recorded from each shank to a particular thalamic nucleus. This was aided by calretinin immunohistochemistry, which provided clear boundaries between rmPulv/cmPulv (calretinin+ cell bodies), lPulv (calretinin-), and dLGN (calretinin+ axons). While calretinin does not distinguish between rmPulv and cmPulv, the distinction was clear because V1 axons were only in rmPulv, but not cmPulv (Zhou et al., 2017). Thus, for pulvinar recordings, we only included units in our analyses that were recorded on shanks that passed through fluorescent V1 axons (in the absence of V1 axons, e.g., L5ET-dtA experiments, we compared our calretinin staining and the shape of the dentate gyrus to the Allen Brain Institute’s Reference Atlas and our own images with both calretinin staining and V1 axons). Because the boundaries of dLGN were perfectly clear with calretinin staining, any shanks in the dLGN, even if they did not pass through L6CT terminals, were included in our analyses. The dorsal/ventral boundaries of our thalamic recordings were determined by assessing visual responsivity (see below for criteria for being “visually responsive”); since the top channels of our recording probe were typically in the hippocampus and thus virtually silent, we considered the most dorsal channel with a visually responsive unit as the top of dLGN/pulvinar (hence, “depth” in LMI plots throughout this study are relative to the position of this first channel). Similarly, the ventral boundary of the thalamus was identified as the last channel with a visually responsive unit, and all units recorded from those and intervening channels were included in analyses.

Units were considered “visually responsive” if there was a significant difference in spike counts during either the initial (first 200ms) or sustained (250-1750ms) period following the onset of the drifting grating stimulus, compared between preferred visual stimulus trials (i.e., the direction with the biggest difference from baseline) and blank trials (using the Wilcoxon rank-sum test with p=0.025 significance threshold) in which the mouse was stationary. For non-optogenetic experiments (i.e., diphtheria toxin experiments), there were no “blank” trials and so instead the first 200ms or sustained 500ms (250-750ms) periods from visual stimulus onset were compared to the same durations at the start of the trial (pre-stimulus) with the paired Wilcoxon signed-rank test.

A similar approach was used to determine the significance of a unit’s light modulation; spike counts during the sustained (250-1750ms from visual stimulus onset) period were compared between blank (stationary) trials with and without light stimulation. We used the Benjamini-Hochburg method for false discovery rate correction to limit the estimated rate of false positives among our modulated cells to <10% (typical significance threshold ∼p=.025, no less than .01). These significantly modulated cells were then classified as “suppressed” or “enhanced” based on whether their spontaneous light modulation index was less than (suppressed) or greater than 0 (enhanced). In joint SC- and L5ET/V1-inactivation experiments where we sought to separately assess the effects of four different light stimulation conditions (no light, SC-inactivation, L5/V1-inactivation, SC&L5-inactivation), instead of the rank-sum test we used the Kruskal-Wallis test with the Dunn–Šidák post-hoc test. We did this separately for both visual (preferred direction) and blank trials and designated units as SC-suppressed, L5/V1-suppressed, or suppressed by both if comparisons were significant for either their visual or spontaneous activity (since we noticed that unlike with L5/V1-inactivation, SC inactivation often impacted visual but not spontaneous activity (Casanova and Molotchnikoff, 1990)). As an additional catch for false positives, units had to be significantly (p<0.025) suppressed in their visual or spontaneous activity during the combined inactivation condition (L5/V1- and SC-inactivation) as well as in at least one other inactivation condition to be classified as SC-suppressed, L5/V1-suppressed, or suppressed by both.

For V1, the short recordings of cortical activity in response to screen flashes (see Visual Stimulation above) were used for current source density (CSD) analysis to determine recorded units’ laminar position. We used CSDPlotter (Pettersen et al., 2006) to compute the second spatial derivative of the low-pass-filtered (<1000Hz) local field potential during the transitions between screen-off and screen-on periods. We then assigned electrode channels to cortical layers based on typical spatiotemporal patterns of current sources and sinks immediately following a screen flash (see Figure 2C), and units whose largest voltage deflection was picked up from a particular channel were assigned to that channel’s layer. Units were classified as “fast-spiking” if the time from the trough to the peak of their waveforms was less than .475ms (which typically marked a clear division in the bimodal distribution of trough-to-peak times); all others were considered “regular-spiking”. Units were considered putatively opsin-expressing (“inactivated” units) if they were regular-spiking, in the expected layer (L5 or L6 in Npr3-Cre and Ntsr1-Cre mice, respectively), their visually evoked activity was significantly light modulated (as described above), and LMI_vis_ <-.33 (i.e., >50% suppression of visually evoked activity).

#### Quantification and analysis – drifting gratings experiments

Average visually evoked and spontaneous firing rates (FR_vis_ and FR_spont_) were calculated in the window 250-1750ms after stimulus onset across visual and blank trials, separately for trials in different light stimulation conditions. Only “stationary” (speed < 2cm/s) trials were used for these calculations because running has been shown to affect firing rates in V1 (Niell and Stryker, 2010) and the thalamus (Erisken et al., 2014), and our mice did not run very often (typically <10% of trials). Light modulation indices (LMIs) were then calculated as the difference divided by the sum of average firing rates (FR_vis_ or FR_spont_) in no-light versus light stimulation conditions. The visually evoked FR change (Figure 3J) was calculated as the absolute value of the difference between FR_vis_ and FR_spont,_ separately for each light condition. F1 responses were calculated by computing the Fast Fourier Transform (FFT) of the baseline-subtracted (baseline from the same time window in blank trials for each light stimulation condition separately) average peristimulus time histogram (10ms bins) across preferred direction trials during the sustained response period (250-1750ms after visual stimulus onset). The F1 response was then the component of the resulting spike-response amplitude spectrum at the temporal frequency of the visual stimulus (2Hz). Bursting rates were calculated as the ratio of the number of spikes occurring in bursts to the total number of spikes during the sustained stimulation period in visual and stationary trials, separately for each light condition. Burst spikes were defined as those preceded by an inter-spike interval (ISI) ≥ 100ms and followed by an ISI ≤ 4ms (first spikes in a burst), as well as all subsequent spikes which were preceded by an ISI ≤ 4ms.

#### Quantification and analysis – RF mapping experiments

For each unit, we generated a peristimulus time histogram matrix M of average baseline-subtracted firing rates for each unique stimulus type (white or black square at each possible stimulus position under each light stimulation condition; baseline = mean of first 20ms) at each 10ms time bin following the start of the stimulus (250ms total stimulus presentation = 25bins). We then performed singular value decomposition (SVD) of this matrix M:

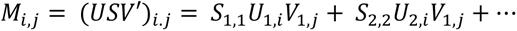

to extract spatial (*U*_1,*i*_) and temporal (*V*_1,*j*_) filter approximations of the spatiotemporal receptive field.

The following criteria were then used for identifying significant receptive fields:

1. The relative variance of the first singular value 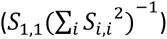 was >10%, AND the peak of the temporal (*V*_1,*j*_) filter exceeded the mean over the first three time bins 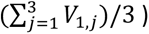 by at least 5 standard deviations.
2. The “spatial SNR” of *U*_1*,i*_ (reshaped to *U*_1*,x,y*_) - the ratio of the maximum variance at any x/y position to the mean variance at the furthest two x/y positions, across x and y dimensions separately - exceeded the 2.5% tails of a null distribution of spatial SNRs (from 1000 shuffles of entries in *U*_1*,x,y*_) in either x or y dimensions. In other words, there was a non-random spatial organization of the estimated receptive field. OR, units also passed this step if a value of *U*_1*,x,y*_ exceeded 5 standard deviations above the mean of *U*_1*,x,y*_ (otherwise, units with very small RFs - most common in dLGN - did not pass the spatial SNR criterion).
3. At least one value of *U*_1*,i*_ was significant relative to a null distribution (from SVD of 500 shuffled *M* matrices), with a significance threshold corrected for <20% FDR (Benjamini-Hochburg method).

For units who passed these criteria for at least 1/4 of their computed receptive field estimates (on and off subfields, light on and light off conditions), we then used the p-values generated in step 3 as a mask for the spatial filter map (*U*_1*,x,y*_). This significant RF map was then bilinearly interpolated to 1° resolution, smoothed with a 2D Gaussian filter (σ=4), and binarized, from which the borders and area of the estimated RF could be determined.

For analyses of retinotopic alignment, we used the estimated receptive field OFF- and ON-fields exclusively from no-light (i.e., control) conditions. RF distance was calculated as the magnitude of the vector connecting the center of all RFs from a single V1 penetration (median x and y positions of all individual unit RF centers) to the center of each thalamic RF, separately for OFF- and ON-subfields. Whichever RF distance (between OFF- and ON-subfields) was shorter was that unit’s final RF distance used for analysis in Figure 4.

For ease of interpretation, all RFs in this manuscript are illustrated as baseline-subtracted firing rates (pre-SVD) at the peak timepoint (i.e., *j* at which *V*_1,*j*_ is maximal). When the peak timepoint differed between light stimulation conditions, the same peak timepoint (from the no-light condition) was used for all conditions.

#### Quantification – cell counting

For quantification of L6CT cells in Ntsr1-Cre/Ai14 mice, we acquired z-stack images at 20x magnification with a Zeiss LSM880 confocal microscope. The counted region was restricted to the area of V1 (as identified by a thicker L4 in the DAPI channel and the relative lack of retrograde labeling in upper L6 relative to more lateral and medial cortical areas, as consistently described in other studies; Blot et al., 2020; Roth et al., 2016; Souza et al., 2020). Within this area, L6 was then split into partitions (as in Frandolig et al., 2019); cells in the upper 10-40% of L6 were counted as L6_upper_, cells in the lower 60-110% were counted as L6_lower_, and all cells throughout the full depth of L6 were included for “all L6” quantification. Only animals in which the CTB injection was confined to the pulvinar and did not spread into dLGN were included in this quantification (n=3). Two or three sections spaced 200-400µm apart were counted per animal using FIJI.

For quantification of L5ETs, we acquired z-stack images at 20x magnification with an Olympus BX63 microscope. For quantification in Npr3-Cre mice, the area of GFP expression (from AAV injection) and the mCherry+ pulvinar axons (dense in L4 of extrastriate areas but not V1) were used to set the counting boundaries in V1. Three sections spaced 50-150µm apart were counted in each of four animals. For quantification of L5ETs with different subcortical targets, cells retrogradely labeled from the SC with CTB-647 were the most spatially restricted and thus were the main determiners of counting boundaries in V1 (V1 boundaries identified by mCherry+ axons as before). Since the SC injection was also the most difficult because of its close proximity to retrosplenial cortex, animals were only included for quantification if all three injections were well targeted and yielded retrograde labeling in V1 but not in the LD thalamic nucleus (a major input to retrosplenial cortex; n=3). Five sections, each separated by 150µm, were counted for each animal. FIJI software was used for all cell counting.

#### Statistical analysis

For paired comparisons, we used the Wilcoxon signed-rank test. For independent comparisons with two groups, we used the Wilcoxon rank-sum test, and for more than two groups we used the Kruskal-Wallis test with the Dunn–Šidák post-hoc test for multiple comparisons. Statistical significance was assessed at the 0.05 level unless otherwise specified.

---

**Figure S1.**
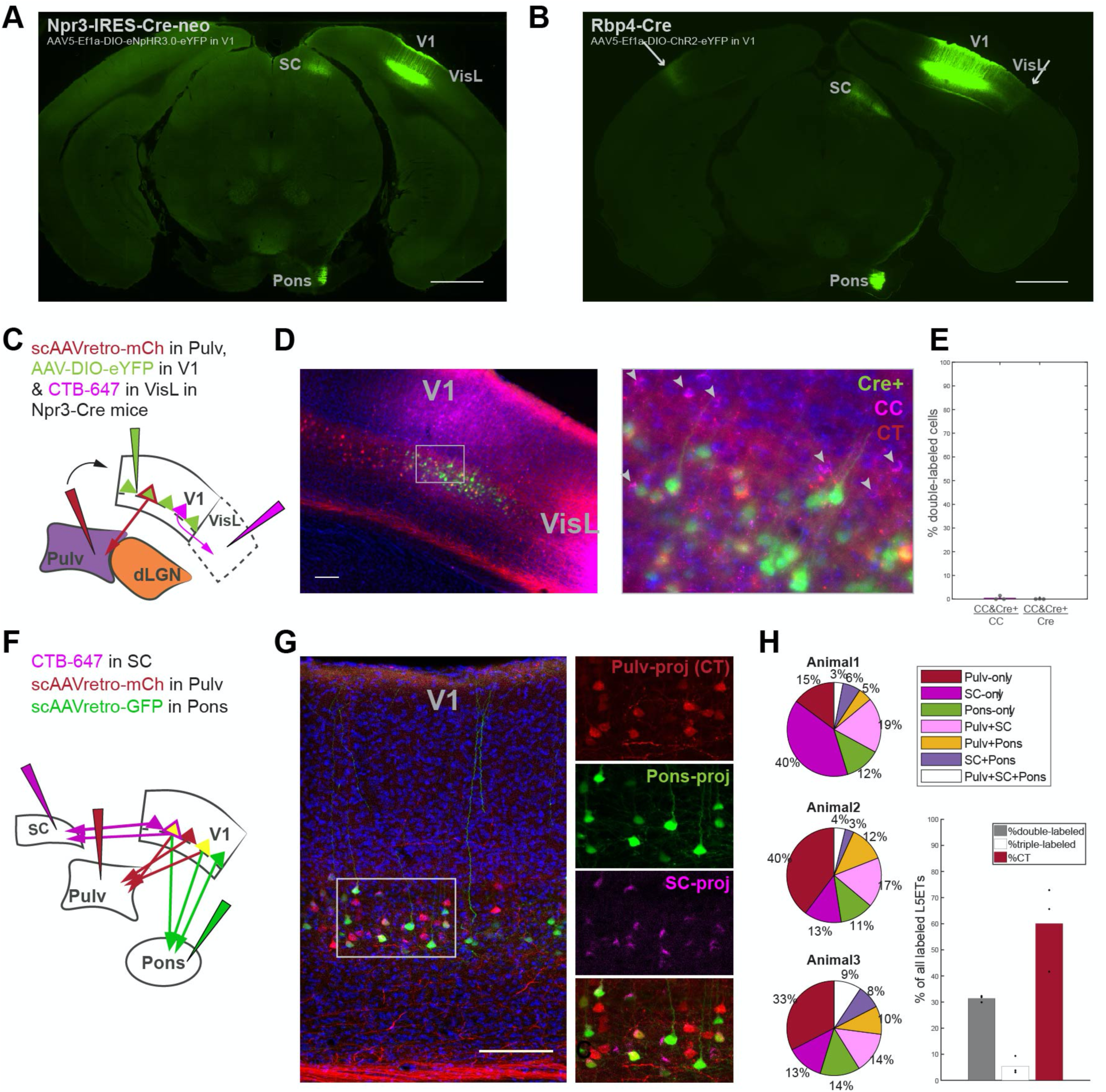
Additional data pertaining to L5 extratelencephalic (L5ET) cells and the Npr3-Cre mouse line, related to Figure 1. (A-B) Epifluorescence images of a Npr3-Cre mouse injected with AAV5-EF1a-DIO-eNpHR3.0-eYFP in V1 (A), and a Rbp4-Cre mouse injected with AAV5-Ef1a-ChR2-eYFP in V1 (B). Axons are visible in subcortical targets SC and pons in both cases, yet cortical axons (in contralateral cortex and in lateral visual cortex (VisL), indicated by arrows) are only visible in Rbp4-Cre mice, even at high exposures. 1mm scale bars. (C) Corticothalamic and CC cells were retrogradely labeled from pulvinar and VisL, respectively, in Npr3-Cre mice injected with an AAV encoding Cre-dependent eYFP in V1 (three out of four of the same experiments depicted in Figure 1F). (D) Epifluorescence images (at 20x, maximum intensity projection) of V1 and VisL. Arrowheads in higher-magnification image (from boxed area) indicate retrogradely labeled CC cells which are neither mCh+ (CT) nor eYFP+ (Cre+). 100µm scale bar. (E) Quantification of double-labeled (CTB+ and eYFP+) cells out of all CC (CTB+) and all Cre+ (eYFP+) cells. Only one doubled-labeled cell was found in any of the three animals, hence values are virtually zero (0.49% and .05%, means across n=3 animals). F) Three different retrograde tracers were injected into different subcortical targets of L5ETs (SC, Pulvinar and Pons) in C57 mice. (G) Confocal image of retrogradely labeled cells in V1, including higher-magnification single- and composite-channel images of boxed region. 100µm scale bar. (H) Proportions of different retrogradely labeled cell types in each of three experimental animals. Bottom-right: proportions of double-labeled (2 subcortical projection targets), triple-labeled (all three projection targets), and pulvinar-projecting (CT) cells across all three mice. Proportions of CT cells are likely overestimates, as we observed more numerous and widespread retrogradely labeled cells from the pulvinar than from either of the other subcortical injections. Bars indicate means across n=3 animals.

**Figure S2.**
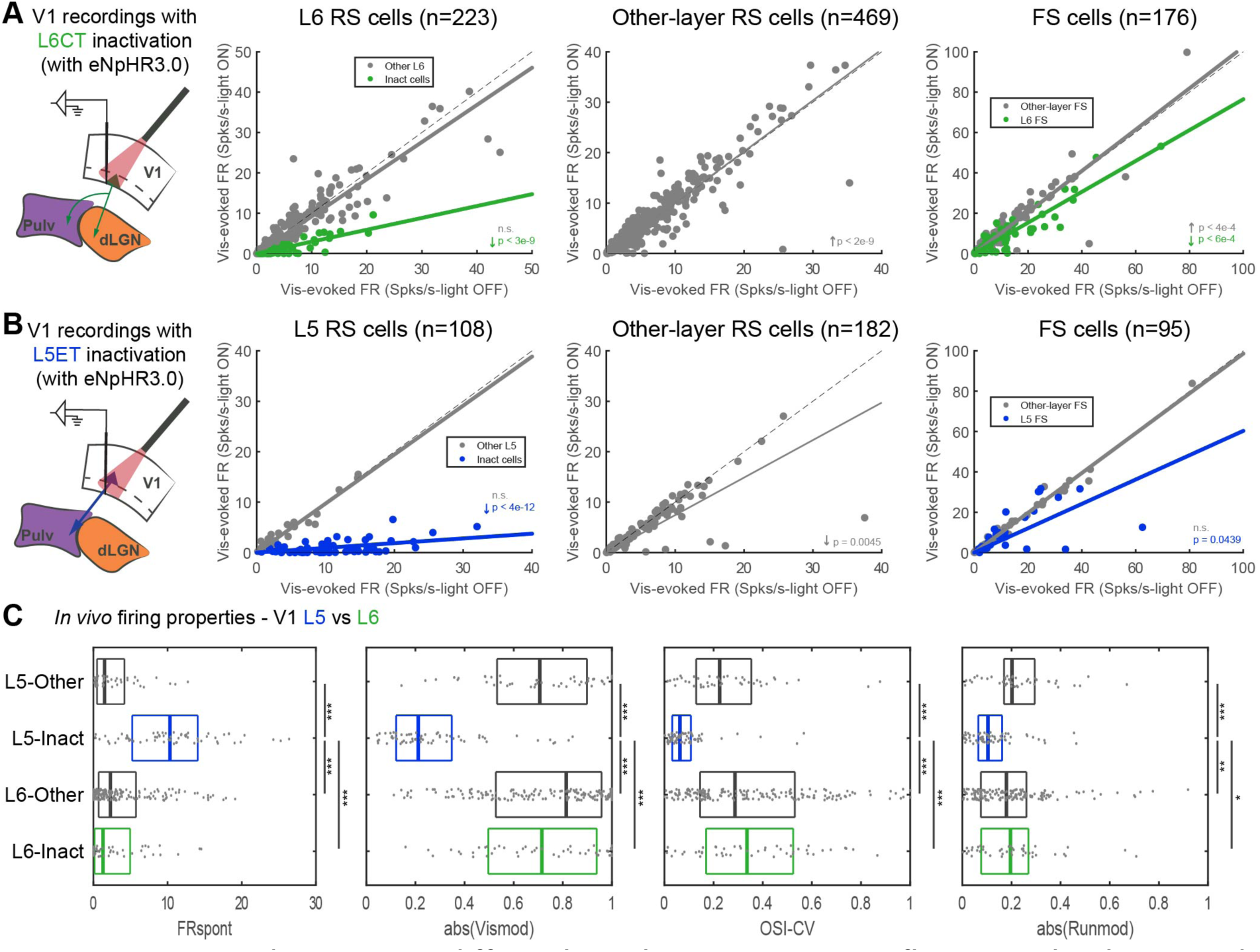
L6CTs and L5ETs in V1 differ in how their inactivation influences other layers and in their firing and tuning properties, related to Figure 2. (A) Data from V1 recordings in Ntsr1-Cre mice injected with Cre-dependent halorhodopsin. Visually evoked firing rates, with versus without L6CT inactivation for (left to right): regular-spiking (RS) cells within L6 (“inactivated” cells or “other”); all other RS cells (layers 2-5); and all fast-spiking (FS) cells (L6 vs. all other layers). Significance values are from Wilcoxon signed-rank tests, and arrows indicate the direction of significant modulation. (B) Same as (A) but for L5ET inactivation experiments in Npr3-Cre mice (comparing L5 versus other layers). (C) Comparison of *in vivo* firing and tuning properties among “other” and “inactivated” L5 RS cells (in Npr3-Cre mice) and L6 RS cells (in Ntsr1-Cre mice). Left to right: boxplots of medians and quartiles of spontaneous firing rates (from blank trials); visual modulation index magnitudes (difference divided by sum of average firing rates from preferred visual and blank trials under no-light conditions); orientation selectivity indices (1-circular variance); and running modulation index magnitudes (difference divided by sum of average visually evoked firing rates from running versus stationary trials, only for experiments with >5% running trials). ***p<0.001, **p<0.01, *p<0.05, Kruskal-Wallis non-parametric test with the Dunn–Šidák post-hoc test for multiple comparisons.

**Figure S3.**
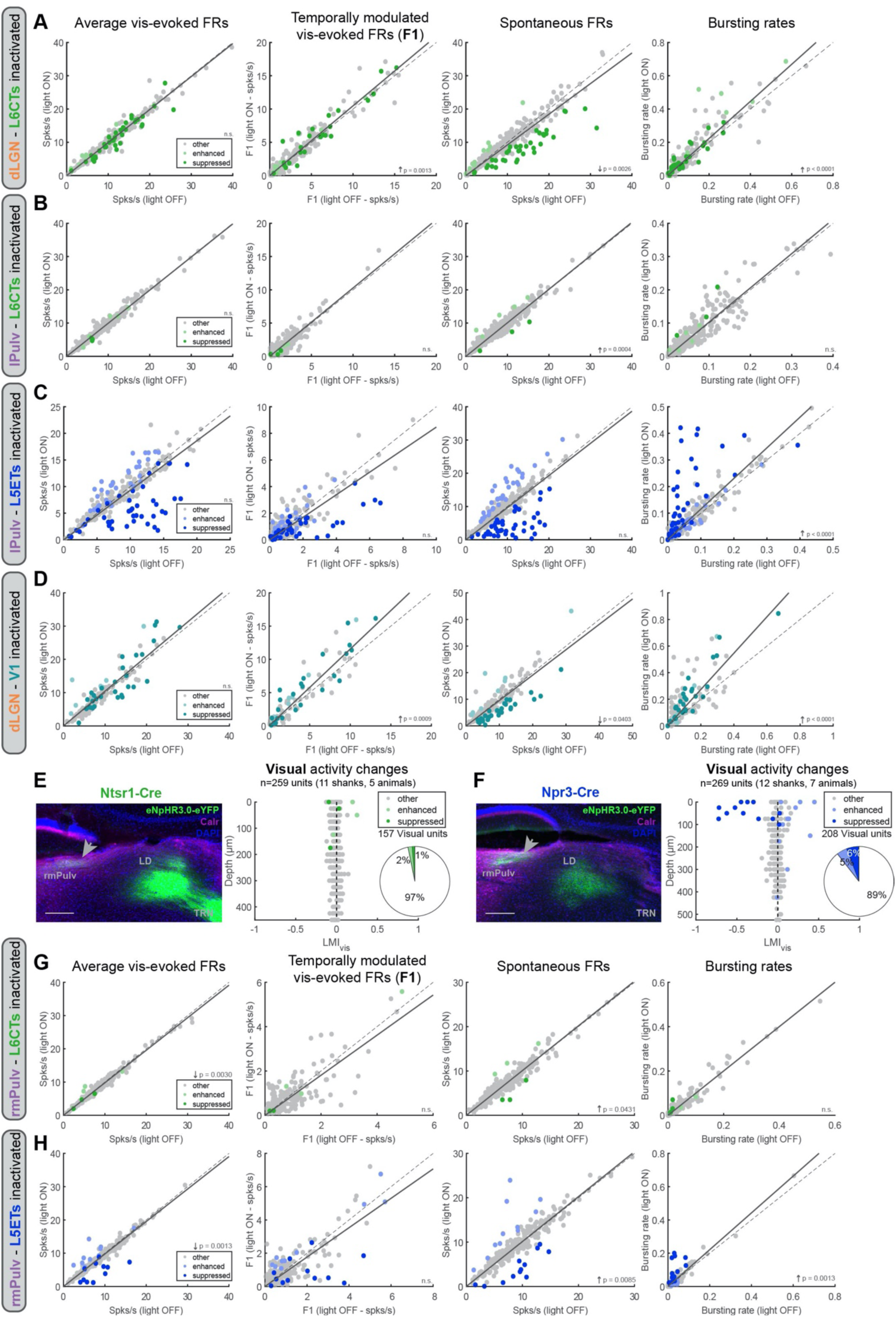
Population-level effects of layer-specific and non-specific V1 inactivation in the dLGN, lateral pulvinar (lPulv) and rostromedial pulvinar (rmPulv), related to Figure 3. (A) Effects of L6CT inactivation on dLGN units’ (left to right): average visually evoked activity across all visual trials; temporally modulated responses (i.e., F1) to the preferred visual stimulus; spontaneous firing rates (from blank trials); and bursting rates. Units are colored according to whether their spontaneous activity was suppressed, activated, or non-modulated (“other”) by light stimulation (same classification as in Figure 3). (B-D) Same as (A) but for: (B) lPulv units recorded with L6CT inactivation; (C) lPulv units recorded with L5ET inactivation; and (D) dLGN units recorded with non-specific V1 inactivation (n=2 PV-Cre/Ai32 mice, n=2 Dlx5-Flp mice with AAV-fDIO-ChR2-eYFP injection in V1; pulvinar data from these experiments are presented in Figures S5 and S7). (E) Left: L6CT axons in the rmPulv, as well as the lateral dorsal nucleus (LD) and thalamic reticular nucleus (TRN) in halorhodopsin-expressing Ntsr1-Cre mice. rmPulv stains positive for calretinin (unlike lPulv). Scale bar = 200µm. Right: Light modulation of visual activity (LMI_vis_) by depth for all rmPulv units in L6CT inactivation experiments. Inset: proportion of units whose spontaneous activity was significantly suppressed, enhanced or unmodulated (same criteria as for Figure 3). (F) Same as (E) but for rmPulv recordings in L5ET inactivation experiments (Npr3-Cre mice). (G-H) Same as (A-D) but for rmPulv units recorded with L6CT inactivation (G) or L5ET inactivation (H). All p-values are from Wilcoxon signed-rank tests (all units combined); arrows indicate direction of significant change.

**Figure S4.**
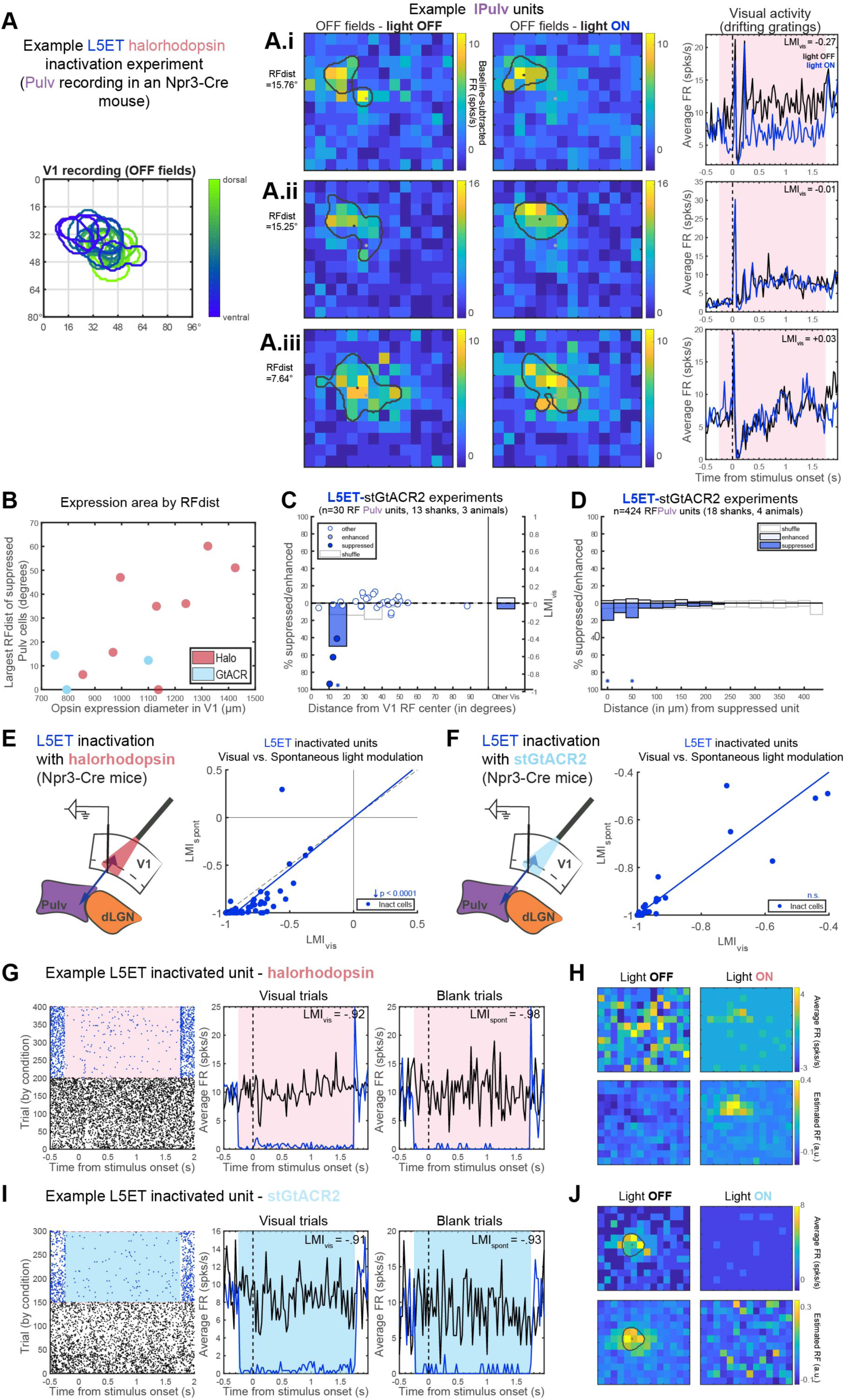
Effects of halorhodopsin and/or stGtACR2 L5ET inactivation on retinotopically aligned pulvinar units and on visual versus spontaneous activity in L5ETs themselves, related to Figures 4 and 5. (A) An example L5ET inactivation experiment (with halorhodopsin) in which both V1 and lPulv was recorded from (not simultaneously). Left: overlayed RFs (in response to luminance decreases, i.e., OFF fields) recorded from V1. (A.i-iii): three example lPulv units’ RFs with and without light for L5ET inactivation (middle), and average PSTHs in response to drifting gratings (right). RF maps depict baseline-subtracted firing rates in response to sparse noise stimuli (luminance decreases) presented at each position in the grid at the timepoint of peak response. Black outlines and dots indicate the boundaries and centers of their estimated RFs (see Methods for details). Grey dots mark the center of V1 RFs (accounting for the wider grid of stimulus positions that was used in the pulvinar than in the V1 recordings). “RFdist” is the distance (in degrees) from each unit’s RF center to the center of all V1 RFs. Note that A.i and A.ii have similar RF locations and distances, yet only A.i is significantly suppressed by L5ET inactivation during drifting grating experiments. Meanwhile, A.iii has an even shorter RF distance to the inactivation/recording site in V1 yet was unaffected by L5ET inactivation. (B) Opsin expression diameter in V1 versus the largest RF distance of a significantly suppressed pulvinar cell recorded in that animal, among halorhodopsin- and stGtACR-expressing animals. (C) Relationship between light modulation (in drifting grating experiments, Figure 5) and RF distance in L5ET-stGtACR2 experiments in which sparse noise stimulation was used for both pulvinar and V1 recordings. Dots indicate LMI_vis_ (right y-axis) of enhanced, suppressed, and other cells (same as Figure 5: classified from blank trials). Bars indicate the proportion of units significantly suppressed or enhanced (left y-axis), binned by retinotopic distance to V1 inactivation site (10° bins). ‘Other Vis’ are all other units from the same experiments whose RFs could not be determined. White bars reflect means of 1000 shuffled distributions (shuffled separately for each experiment). Asterisks indicate where actual proportions fell beyond either tail (2.5%) of shuffled distributions. Fewer significant RFs were identified in these experiments than in halorhodopsin experiments (Figure 4), likely because we used twice the number of light conditions (including SC inactivation, Figure 7) and thus fewer repeated stimulus presentations. (D) In L5ET-stGtACR2 experiments, percent of all unit pairs - consisting of two units, at least one of which was significantly suppressed, recorded on the same shank in the same experiment - in which the second unit was suppressed or enhanced, binned by vertical distance between the channels from which those units were recorded. White bars reflect means of 1000 shuffled distributions (shuffled separately for each recording shank). Asterisks indicate where actual proportions fell beyond either tail (2.5%) of shuffled distributions. (E-F) Inactivation of halorhodopsin-expressing (E) or stGtACR2-expressing (F) L5ET cells in V1 with red (617nm) or blue (455-470nm) LED light, respectively. Right: light modulation indices across visual versus blank trials (LMI_vis_ vs. LMI_spont_) for putatively opsin-expressing L5ETs in V1. Individual units’ LMIs were typically more negative (i.e., stronger inactivation) in blank than in visual trials in halorhodopsin experiments (p<0.0001, Wilcoxon signed-rank test), but not in stGtACR2 experiments. (G) An example regular-spiking L5 cell that is putatively halorhodopsin-expressing given its high degree of inactivation. Left: raster plot. Middle: PSTH of average firing rates across visual trials. Right: PSTH of average firing rates across blank trials. Shading indicates the period of LED stimulation. (H) Top row: the same example unit’s average baseline-subtracted FRs at the peak delay time in response to sparse noise stimulus presentations (luminance increases), without (left) or with (right) LED stimulation for L5ET inactivation. Bottom: the estimated spatial receptive field from singular value decomposition of the matrix of PSTHs across all stimulus locations (see Methods for details). Despite visual and spontaneous LMIs <-0.90, this unit does not have a discernable RF under normal conditions, but with background activity levels suppressed by LED stimulation, the signal-to-noise ratio improves such that a clear RF emerges. (I) An example putative stGtACR2-expressing L5ET unit. (J) Same as (H) but for the unit in (I). In this case, the unit’s response to stimuli presented in its RF is entirely abolished.

**Figure S5.**
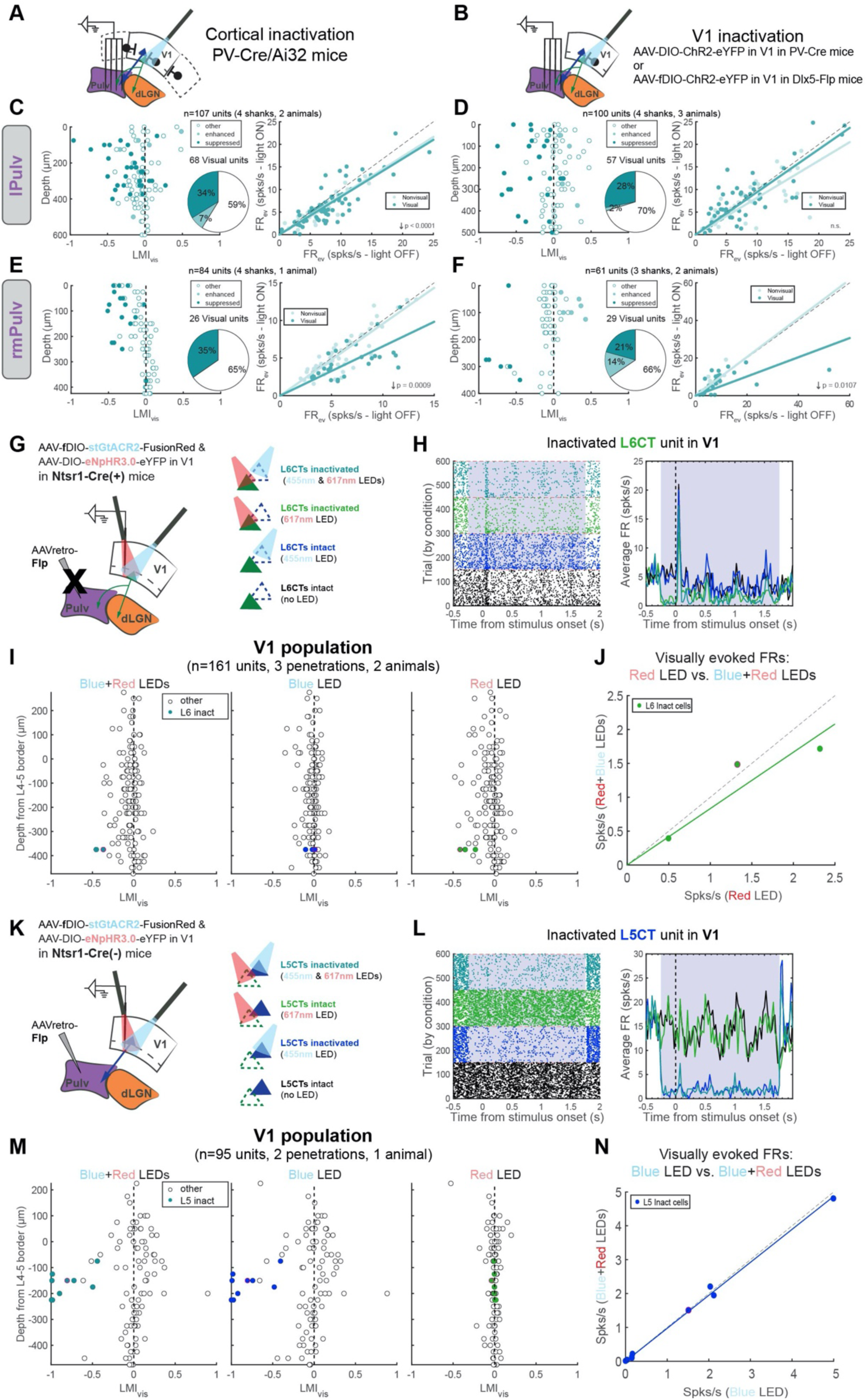
V1 inactivation experiments and dual-wavelength optogenetics control experiments, related to Figure 6. (A-B) Schematic of V1 inactivation experiments in: (A) PV-Cre/Ai32 mice, in which PV interneurons across the brain expressed ChR2 but blue LED stimulation was targeted to V1; and (B) V1 AAV-injected animals (PV-Cre mice with an AAV-DIO-ChR2 injection in V1, n=2; or Dlx5-Flp/CCK-Cre mice with AAV-fDIO-ChR2 injection in V1, n=2). (C-D) lPulv recordings in PV-Cre/Ai32 (C) and AAV-injected (D) mice. Left: LMI_vis_ values by depth for suppressed, enhanced and non-modulated (“other”) units (classified on basis of their spontaneous activity, as in Figures 3 and 5). Inset: pie chart showing proportions of different unit types among all visually responsive units. Right: FR_vis_ in light OFF versus light ON (V1 inactivated) conditions for visual and non-visual cells. P-values are from Wilcoxon signed-rank tests; arrows indicate the directions of significant modulations. (E-F) Same as (C-D) but for units recorded in rmPulv. (G) Schematic of dual-wavelength optogenetics control experiments. Ntsr1-Cre(+) mice were injected with Cre-dependent halorhodopsin and Flp-dependent stGtACR2 (but AAVretro-Flp was NOT injected to pulvinar, so only halorhodopsin was expressed in L6CTs). Right: four different LED stimulation conditions (randomly interspersed). (H) Example L6CT inactivated unit in V1. Left: raster plot, with trials organized by LED stimulation condition. Right: PSTH of average firing rates across visual trials. Grey shading indicates the period of LED stimulation. (I) Visual light modulation (LMI_vis_) calculated from trials with both LEDs on (left), blue LED only (middle), and red LED only (right) trials. Colored units are those whose FRs were significantly suppressed by at least 50% in visual trials with both LEDs, and the dot outlined in pink is the example unit in (H). (J) Visually evoked firing rates of L6 inactivated cells in conditions with the red LED alone compared to both LEDs together. Their FRs were largely unaffected by blue LED stimulation. (K) Same as (G) except AAVretro-Flp was injected to the pulvinar, and injections were made in Ntsr1-Cre(-) mice, resulting in only L5CT expression of fDIO-stGtACR2. (L) Example L5CT inactivated unit in V1. (M) Same as (I) but in a L5CT inactivation experiment, and colored dots are L5 inactivated units (outlined dot is the example unit in (L)). (N) Visually evoked firing rates of L5 inactivated cells in conditions with the blue LED alone compared to both LEDs together. FRs were unaffected by red LED stimulation.

**Figure S6.**
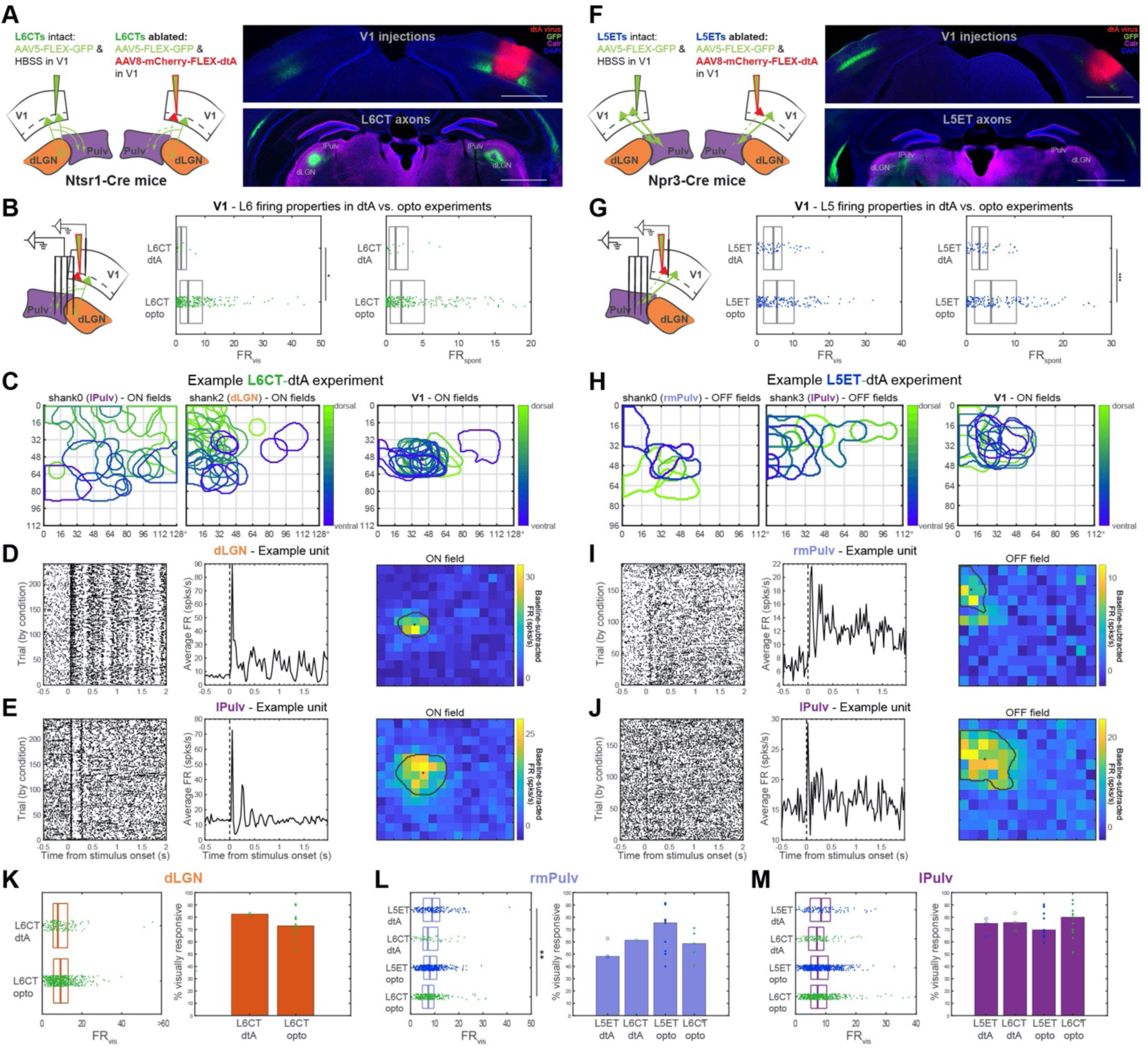
Targeted ablation of V1 L5ETs or L6CTs does not impair visual activity in the visual thalamus, related to Figures 5 and 6. (A) Experimental design of L6CT ablation. Left: in one hemisphere of Ntsr1-Cre mice, two AAVs - one encoding Cre-dependent GFP and the other encoding Cre-dependent diphtheria toxin (dtA) with non-Cre-dependent mCherry were injected into V1. This resulted in no L6CT GFP expression anywhere that the dtA virus spread to, while the GFP virus spread further (due to the difference in AAV serotypes) and labeled L6CTs on the outskirts of the ablation zone. In V1 of the opposite hemisphere, the same Cre-dependent GFP virus was injected without the dtA virus (but with an equal volume of HBSS to attain the same dilution). Right: images of coronal sections containing V1 injection areas (top) and sections containing GFP+ axons in the visual thalamus (bottom). Notice that only in the dtA-injected hemisphere is there a considerable gap in GFP+ axons in the dLGN, indicating that L6CTs were successfully ablated. Scale bars = 1mm. (B) Simultaneous V1 and thalamic recordings in L6CT-ablated mice. Visually evoked (middle) and spontaneous (right) firing rates of regular-spiking L6 V1 units in dtA experiments are depicted and compared to optogenetics experiments (L6 units from halorhodopsin and stGtACR2 experiments are combined). Boxplots depict medians and quartiles. There was a significant reduction in L6 visually evoked firing rates (p=0.0132), but not spontaneous firing rates (p>0.3) in dtA ablation experiments (Wilcoxon rank-sum tests). (C) Overlayed RFs of simultaneous dLGN, lPulv, and V1 recordings from an example experiment. (D-E) Example dLGN (D) and lPulv (E) units recorded in the same experiment as in (C). These units exhibited robust visual responses to drifting gratings (left, raster plot; middle, PSTH of average FRs across all visual trials), even though they were retinotopically aligned with the recorded area of L6CT ablation (right; compare to V1 RFs in (C)). RF maps depict mean baseline-subtracted FRs for each ON stimulus position at the timepoint of peak response. (F) L5ET ablation experimental design - same as (A) but in Npr3-Cre mice. (G) Same as (B) but for L5 units in dtA versus optogenetic experiments. L5 spontaneous firing rates (p=0.0005), but not visually evoked firing rates (p>0.1), were significantly reduced in dtA experiments (Wilcoxon rank-sum tests), indicating successful L5ET ablation (because these cells have uniquely high spontaneous firing rates, Figure S2C). (H) Overlayed RFs from rmPulv, lPulv and V1, recorded simultaneously in the same example experiment. (I-J) Same as (D-E) but for example rmPulv (I) and lPulv (J) units recorded in the same L5ET dtA experiment as in (H) whose RFs were aligned to the area of L5ET ablation yet maintained their visual responses. (K) Left: Visually evoked firing rates of dLGN units recorded in L6CT dtA experiments compared to L6CT optogenetic inactivation experiments (halorhodopsin and stGtACR2 experiments combined). Dots are individual units. Right: percent of recorded dLGN units which were visually responsive in dtA versus optogenetic experiments. Circles/dots indicate individual animals. (L-M) same as (K) but for rmPulv (L) and lPulv (M) recordings, comparing both L5ET and L6CT dtA and optogenetic inactivation experiments. ***p<0.001, Kruskal-Wallis non-parametric tests with the Dunn–Šidák post-hoc test for multiple comparisons. No other comparisons were significant.

**Figure S7.**
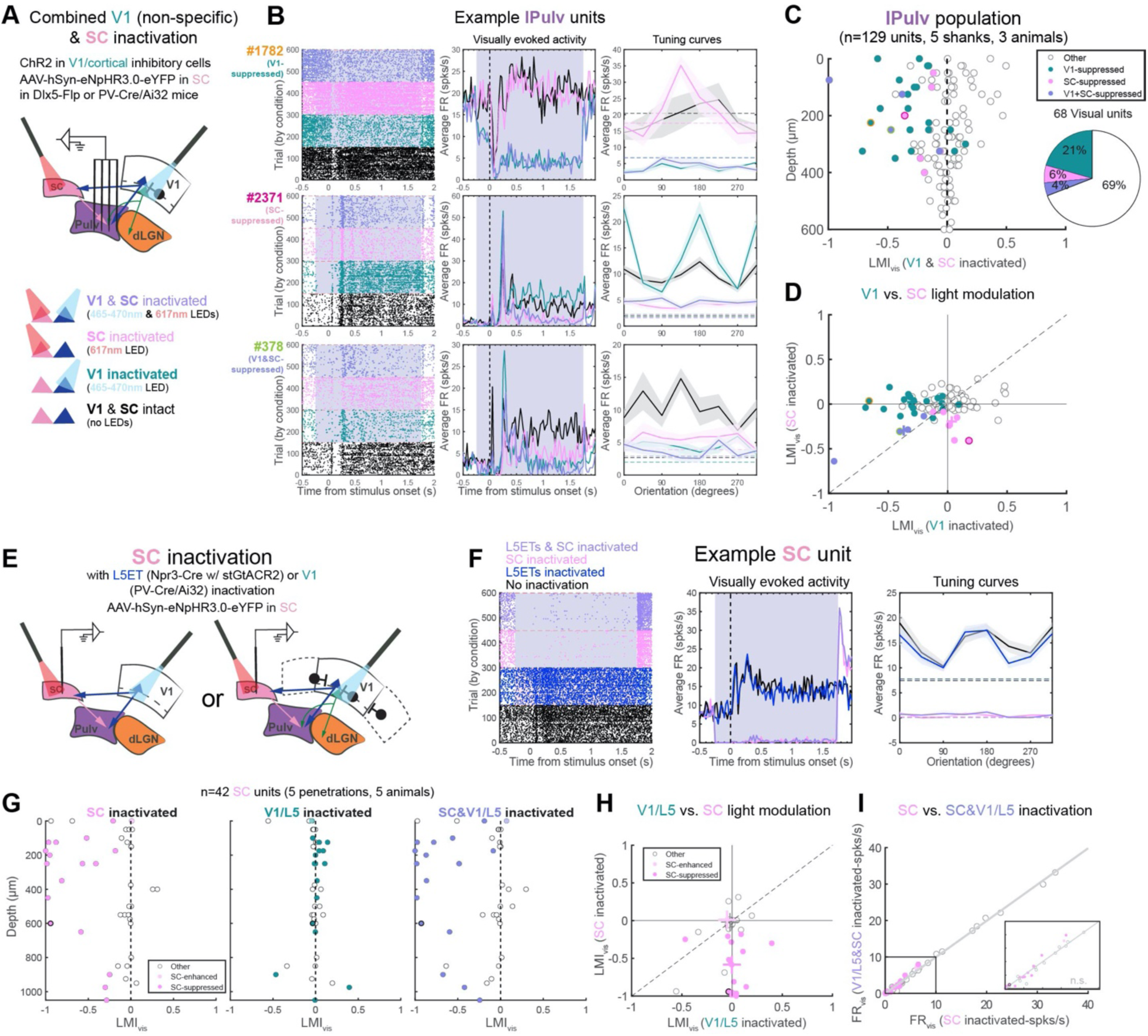
Effects of V1 and superior colliculus (SC) inactivation in the lateral pulvinar (lPulv) and in the SC, related to Figure 7. (A) In vivo pulvinar recordings with dual-optogenetic inactivation of SC and V1 in Dlx5-Flp mice with AAV-fDIO-ChR2 in V1 (n=2) or in PV-Cre/Ai32 mice (n=1). Bottom: Different LED stimulation conditions (same experimental setup as in Figure 7). (B) Three example lPulv units which were suppressed by V1 inactivation (top), SC inactivation (middle; but an increase in activity and tuning with V1 inactivated), or both (bottom). From left to right: raster plots of trials organized by LED stimulation conditions; PSTHs of average FRs across visual trials; and average FRs at each of eight drifting grating orientations under different LED stimulation conditions. Dotted lines in tuning curve plots indicate spontaneous FRs under different inactivation conditions. (C) Light modulation of visually evoked activity (LMI_vis_) from combined V1 & SC inactivation trials, by depth. Units are classified according to whether they were also significantly suppressed by SC inactivation, V1 inactivation, or both (see Methods for details). Color-outlined points correspond to example units in (B). Right: proportion of different suppressed unit classes out of all visually responsive units. (D) Units’ visual light modulation (LMI_vis_) in V1 inactivation versus by SC inactivation trials. Units are colored as in (C). Units which fall in the lower left quadrant were inactivated by both V1 and SC inactivation. Crosses indicate medians of each unit class. (E) Schematic of SC recordings with non-specific halorhodopsin expression for SC inactivation. In addition, either L5ETs specifically (AAV-SIO-stGTACR2 injection in Npr3-Cre mice, as in Figure 7; n=3) or V1 non-specifically (in PV-Cre/Ai32 mice, as above; n=2) were inactivated with a blue LED. (F) An example unit recorded in the SC that was completely silenced by SC inactivation alone or together with L5ET inactivation, but not by L5ET inactivation alone. Same plot descriptions as for (B). (G) Light modulation indices of visually evoked activity (LMI_vis_), calculated from trials with SC inactivation only (left), V1/L5 inactivation only (middle), or combined SC and V1/L5 inactivation (right), by recording depth. Colored dots indicate units whose spontaneous activity was significantly suppressed, enhanced, or unmodulated by SC inactivation alone. The outlined dot is the example unit from (B). (H) Units’ LMI_vis_ in V1/L5 inactivation versus SC-only inactivation conditions. Units are colored as in (G). Crosses indicate medians of each unit class. (I) Visually evoked firing rates with SC inactivation alone compared to combined V1/L5 and SC inactivation. Inset: zoom of boxed region. Units are colored as in (G-H). p’s>0.8 for suppressed cells only or all cells combined; Wilcoxon signed-rank tests.

## Notes

### Competing Interest Statement

The authors have declared no competing interest.

